# Auxin-Induced Actin Cytoskeleton Rearrangements Require AUX1

**DOI:** 10.1101/630798

**Authors:** Ruthie S. Arieti, Christopher J. Staiger

## Abstract

The actin cytoskeleton is required for cell expansion and is implicated in cellular responses to the plant growth hormone auxin. However, the molecular and cellular mechanisms that coordinate auxin signaling, cytoskeletal remodeling, and cell expansion are poorly understood. Previous studies have examined actin cytoskeleton responses to long-term auxin treatment, but plants respond to auxin over short timeframes, and growth changes within minutes of exposure to the hormone. To correlate actin arrays with degree of cell expansion, we used quantitative imaging tools to establish a baseline of actin organization, as well as of individual filament behaviors in root epidermal cells under control conditions and after treatment with a known inhibitor of root growth, the auxin indole-3-acetic acid (IAA). We found that cell length was highly predictive of actin array in control roots, and that short-term IAA treatment stimulated denser, more longitudinal, and more parallel arrays by inducing filament unbundling within minutes. By demonstrating that actin filaments were more “organized” after a treatment that stopped elongation, we show there is no direct relationship between actin organization and cell expansion and refute the hypothesis that “more organized” actin universally correlates with more rapidly growing root cells. The plasma membrane-bound auxin transporter AUXIN RESISTANT 1 (AUX1) has previously been shown necessary for archetypal short-term root growth inhibition in the presence of IAA. Although AUX1 was not previously suspected of being upstream of cytoskeletal responses to IAA, we used *aux1* mutants to demonstrate that AUX1 is necessary for the full complement of actin rearrangements in response to auxin, and that cytoplasmic auxin in the form of NAA is sufficient to stimulate a partial actin response. Together, these results are the first to quantitate actin cytoskeleton response to short-term auxin treatments and demonstrate that AUX1 is necessary for short-term actin remodeling.

**One sentence summary:** The Arabidopsis AUX1 auxin transport protein is necessary for actin cytoskeleton reorganization in response to phytohormone treatment.

## INTRODUCTION

Despite human dependence on plants for food, fiber, and fuel, we do not fully understand the molecular mechanisms controlling plant growth. Many types of plant cells begin life as roughly isotropic but, during development, the cell establishes polar growth where deposition of cell wall materials is restricted to specific axes of the cell, or expansion is anisotropic, allowing the production of mature cells with a myriad of final shapes and sizes. Turgor pressure drives expansion within the confines of cell wall flexibility: certain areas of the plant cell wall are more flexible than others, and are therefore more susceptible to turgor pressure exerted by the vacuole (Szymanski and Cosgrove, 2009; Guerriero et al., 2014). Vesicles are incorporated into certain areas of the plasma membrane and deposit new cell wall material, increasing the cell’s surface area and conducting the cell to grow into specific shapes. Vesicle delivery and exocytosis of vesicle contents of wall materials depend on the actin cytoskeleton (Ketelaar et al., 2003; Hussey et al., 2006; Leucci et al., 2007; Zhang et al., 2019). When actin is disrupted with pharmacological treatments, cells elongate more slowly (Baluška et al., 2001), implicating actin as a crucial player in cell expansion. Although the actin cytoskeleton is required for plant cell expansion (Baluška et al., 2001; Gilliland et al., 2003; Mathur, 2004; Hussey, 2006; Rahman et al., 2007; Kandasamy et al., 2009; Yang et al., 2011; Guerriero et al., 2014), actin’s function in this process is not well understood. Actin is accepted to provide tracks for vesicle delivery (Mathur, 2004; Hussey et al., 2006), but connections have also been made between certain actin arrays and plant growth (ex., Nick et al., 2009; Higaki et al., 2010a; Smertenko et al., 2010; Dyachok et al., 2011; Yang et al., 2011, Yanagisawa et al., 2015), resulting in various hypotheses about actin’s role and/or the significance of specific actin arrays, each with a degree of supporting evidence, much of it circumstantial (Li et al., 2015a; Szymanski and Staiger, 2017).

Actin arrays form an apparently “organized” orientation, with actin bundles roughly parallel to the longitudinal axis of the cell in rapidly growing root epidermal cells in the light (Dyachok et al., 2011). In the dark, where cell expansion is substantially slower, actin exhibits what appears to be a state of “disorganization”: filaments are substantially less aligned relative to the longitudinal axis of root cells (Dyachok et al., 2011). However, data substantiating cause-and-effect are missing from the literature. Whether a longitudinal array is necessary for, coincides with, promotes, or (conversely) is the product of, cell expansion—or whether the “disorganized” array inhibits or coincides with a cessation of expansion—is not understood and is largely unexamined.

In addition to longitudinal actin orientation, various actin arrays have been correlated with cell length or cell expansion. However, there does not seem to be consensus on whether more longitudinal bundles inhibit (Gilliland et al., 2003; Holweg et al., 2004; Rahman et al., 2007) or stimulate (Kandasamy et al., 2009; Yang et al., 2011; Li et al., 2014a) axial cell expansion and tissue growth. Many previous studies linking specific actin organizations with growth or growth inhibition are based on actin or actin-binding protein mutant phenotypes (Gilliland et al., 2003, Kandasamy et al., 2009, Yang et al., 2011; Li et al., 2014a). Others are based on actin responses to drug or hormone treatments (Holweg et al., 2004; Rahman et al., 2007). Therefore, some of the reported actin–cell growth models may be generalized from what might in fact be more discrete responses: cytoskeletal response to a specific external stimulus (drug or hormone) that affects growth via downstream mechanisms; or filament array changes due to an actin-binding protein whose role could be in only one of many aspects of growth.

In fact, what tasks, exactly, actin undertakes during cell expansion and how these tasks drive or participate in expansion are unclear. Bundles potentially inhibit growth by inhibiting transport of growth hormone-related proteins (Nick, 2010). On the other hand, long actin bundles presumably stimulate growth because they provide tracks for vesicle delivery (Szymanski and Cosgrove, 2009; Thomas, 2012). Actin bundles could play a role in regulating osmotic pressure in the vacuole by altering turgor pressure (Higaki et al., 2010a,b; 2011), the main driver of plant cell expansion (Szymanski and Cosgrove, 2009). A recent paper shows that auxin, a known modulator of plant growth that has opposite effects on root or shoot growth (inhibition and stimulation, respectively), constricts vacuolar shape in long-term treatments (6+ h) on root cells, and does so by inducing altered actin arrays (Scheuring et al., 2016). Although this work describes the long-term effects of auxin on actin (Scheuring et al., 2016), what connects short-term auxin treatments with actin rearrangements is not understood. Interactions between auxin signaling pathways and actin are abundant in the literature (reviewed in Zhu and Geisler, 2015), but the mechanics of how the hormone affects the cytoskeleton on a timescale of minutes, and how these interactions stimulate or inhibit growth, are largely unknown.

The molecular players that connect the actin cytoskeleton to auxin perception during short-term responses are unidentified. Auxin reception by AUXIN BINDING PROTEIN 1 (ABP1) was previously suspected to be upstream of cytoskeletal changes in both roots (Chen et al., 2012; Lin et al., 2012) and epidermal pavement cells (Xu et al., 2010; Nagawa et al., 2012; Xu et al., 2014); however, recent works demonstrate that a CRISPR *abp1-c1* mutant exhibited root growth inhibition in the presence of both the known root growth inhibitor, the auxin indole-3-acetic acid (IAA) and the membrane permeable auxin 1-naphthylacetic acid (NAA), just like wildtype plants, indicating that ABP1 likely does not play a significant role in auxin signaling (Dai et al., 2015; Gao, et al., 2015). AUXIN RESISTANT 1 (AUX1) is a plasma membrane-bound auxin/H^+^ symporter in the Amino acid/auxin permease (AAAP) family that is ubiquitous among Eukaryotes. AUX1 appears to be present in all plants as well as some algae, indicating that the protein likely evolved before land plants (reviewed in Swarup and Péret, 2012). Unlike wildtype, *aux1* plants grow in the presence of, IAA but undergo growth inhibition by NAA (Marchant et al., 1999), and AUX1 binds both IAA and NAA with high affinity (Yang et al., 2006; Carrier et al., 2008) and is responsible for 80% of IAA uptake by root hairs (Dindas et al., 2018). AUX1 contributes to short-term, auxin-induced increases in cytosolic H^+^ and, together with the intracellular auxin receptor complex SCF^TIR1/AFB^, increases in cytosolic Ca^2+^ (Dindas et al., 2018). The auxin molecule itself is the signal that SCF^TIR1/AFB^ perceives (Dharmasiri et al., 2005a,b; Kepinski and Leyser, 2005), driving both rapid increases in Ca^2+^ (Dindas et al. 2018) and transcriptional reprogramming (Ulmasov et al., 1999).

To correlate actin arrays with degree of cell expansion, we used quantitative tools to establish a baseline of actin architecture and orientation and individual filament behaviors in root epidermal cells under control circumstances. By plotting measurements of each cell’s actin array against its length, we found that cell length was highly predictive of actin array. We then used acute treatments with IAA to determine the actin response in presumed non- or very slow-growing cells and documented the first short-term actin responses to these growth-inhibitory doses of IAA. Upon analyzing the actin arrays in two *aux1* alleles (the T-DNA insertion mutant *aux1-100* and the null point mutant *aux1-22*), we found that actin failed to reorganize in response to IAA and actin reorganization was only partially restored by NAA. Our data substantiate that AUX1 and cytosolic auxin play a significant role upstream of actin reorganization in auxin signaling.

## RESULTS

### Actin Organization Correlates with Cell Length

Actin organization in living epidermal cells of the root elongation zone, examined with variable angle epifluorescence microscopy (VAEM), displayed a consistent pattern of organization (Baluška et al., 1997; Figure 1A, Supplemental Figure 1A). The actin array in thin, rectangular cells closest to the root apex—the root cap—comprised haphazardly arranged bundles. About 200 µm from the apex, in short, square cells emerging from under the root cap, there appeared to be a marked increase in the abundance of actin filaments, with fewer bundles. Array organization appeared to become gradually more bundled, longitudinal, and sparse as cells increased in length, until reaching the end of the root elongation zone (a demarcation indicated by the first visible root hair initiations). Although this pattern has been observed previously (Baluška et al., 1997; Baluška and Mancuso, 2013), we wondered whether there were quantitative differences in actin organization that could be correlated with cell size, and, potentially, with developmental stage. After plant cells are generated in the root meristem, they spend approximately 4 d progressing through the meristematic region (including the root transition zone) before progressing to the zone of rapid elongation, where they spend mere hours (Beemster and Baskin, 1998, 2000; van der Weele et al., 2003). The consistent progression of aging, growing cells allows comparison and quantification of actin arrays in cells in both the slower-growing late meristematic/transition zone and the zone of rapid elongation. Whether actin bundles inhibit (Gilliland et al., 2003; Holweg et al., 2004; Rahman et al., 2007) or promote (Kandasamy et al., 2009; Yang et al., 2011; Li et al., 2014a) cell expansion remains controversial. We hoped to gain insight into the role of actin bundling in expansion of root epidermal cells, since it is the root epidermis that drives cell expansion in all root layers (Savaldi-Goldstein et al., 2007 [shoots]; Hacham et al., 2011).

**Figure 1.**
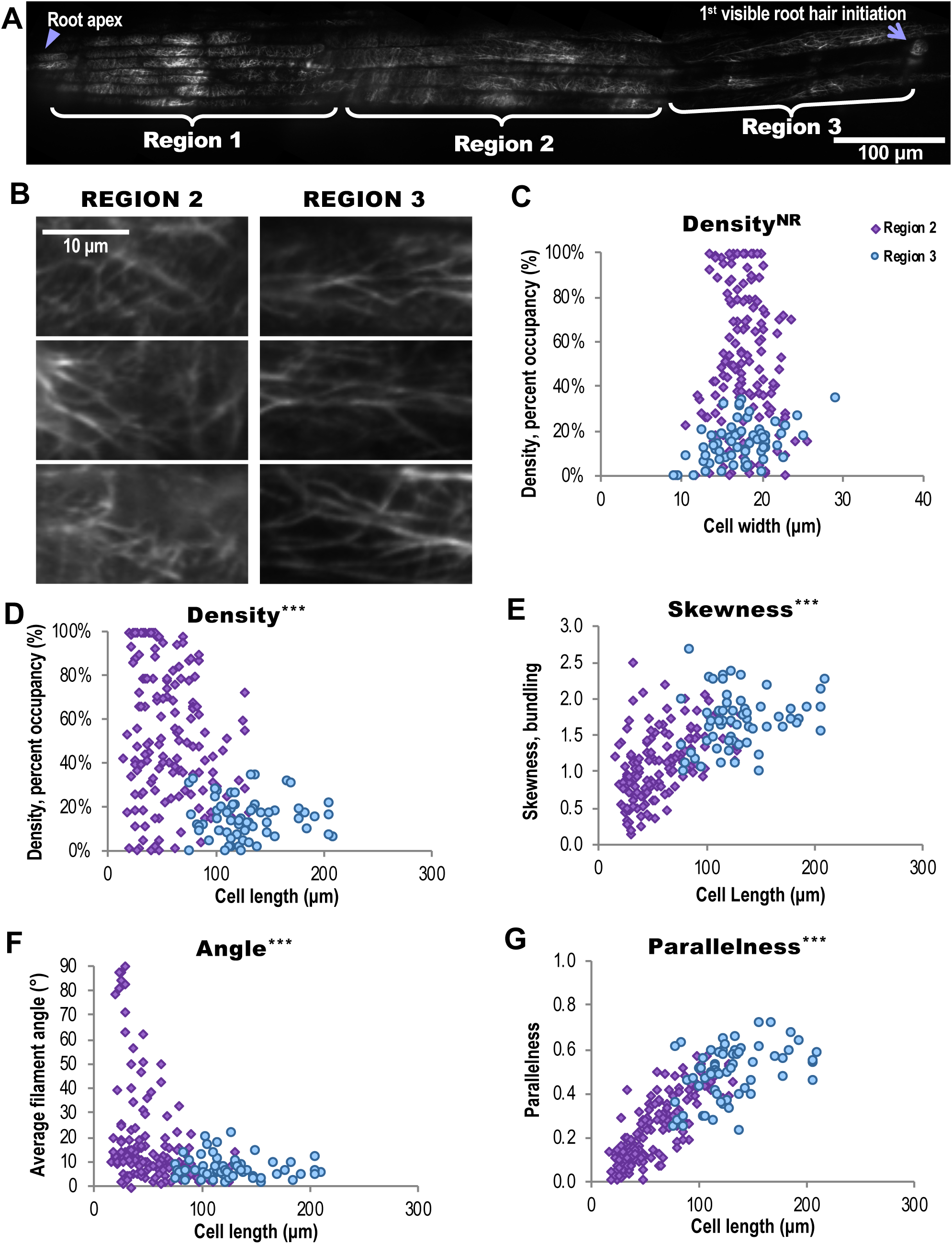
Actin Architecture is Predictive of Epidermal Cell Length in the Root Elongation Zone. **(A)** Mosaic of root elongation zone in an Arabidopsis seedling expressing GFP-fABD2 imaged with variable angle epifluorescence microscopy (VAEM). Arrowhead, root apex; arrow, first root hair initiation. MosaicJ was used to compile 13 original VAEM images. Scale bar, 100 μm. **(B)** Representative images of actin organization in two root regions. Scale bar, 10 μm. **(C)** to **(G)** Quantification of actin architecture or orientation metrics plotted with respect to corresponding epidermal cell length **(D)**, **(E)**, **(F)**, and **(G)** or cell width **(C)** in two regions within the root elongation zone. Filament architecture and orientation were not predictable based on cell width but were highly correlated with cell length. Supplemental Figure 2 shows results for skewness, angle, and parallelness vs. cell width, which also showed no relationship, and Supplemental Figure 1 shows comparisons of Region 2 and Region 3 mean measurements. Mean cell length, Region 2 = 57 ± 28 μm. Mean cell length, Region 3 = 128 ± 34 μm. Region 2 measurements are shown in purple diamonds; Region 3 in blue circles. N = 60–120 cells per region from 20 roots. NR, no predictive relationship; ***, p ≤ 0.0001, Bivariate fit/ANOVA for all data points for each parameter. Results are from one experiment.

To test quantitatively whether actin array organization varies over the course of the root, we took overlapping VAEM images of GFP-fABD2–labeled (green fluorescent protein fused to the second actin-binding domain of Arabidopsis FIMBRIN1) actin filaments from the root apex through the end of the elongation zone. The demarcation between the root cap (here called “Region 1”) and what appeared to be the visible transition zone (here called “Region 2”) was drastic. Isotropic cells that delineate the late meristematic/early transition zone clearly emerge from under the rectangular cells of the presumed root cap. The distinction between Region 2 and what we call “Region 3” was not nearly so definitive as between Region 1 and Region 2, so we first delineated Region 3 by an observable decrease in actin filament abundance (admittedly, a subjective criterion). Representative images showed conspicuous differences in actin arrays in Region 2 and Region 3 (Figure 1B; Region 1, i.e., the root cap, in Supplemental Figure 1A). Aspects of actin organization were quantified as described previously (Higaki et al., 2010b; Ueda et al., 2010; Henty et al., 2011; Li et al., 2012; Cai et al, 2014; Cao et al., 2016). Parameters measured include: percent occupancy or density, the extent of bundling of actin filaments (measured as “skewness” of pixel intensity distribution in an image), parallelness of filaments to each other, and average filament angle relative to a cell’s longitudinal axis (Higaki et al., 2010b; Ueda et al., 2010; Henty et al., 2011; Li et al., 2012; Cai et al, 2014; Cao et al., 2016). These quantitative analyses showed (Supplemental Figure 1B–E) the cortical actin array in root Region 2 was significantly more dense and less bundled than Region 3 (longer cells closer to the first visible root hair initiations). Region 1 was similar in density to Region 2, but more bundled. The filaments and bundles in cells of Region 3 were substantially more longitudinal than those in Region 1 or Region 2. Since cells of the root cap do not follow in the same cell files as Regions 2 and 3 and do not follow the same cell expansion gradient, and since we sought to learn about differences in actin organization over the course of cell expansion, we eliminated Region 1 from further analysis.

Because of the substantial differences in actin organization among epidermal cells within the elongation zone, we hypothesized that if certain actin arrays correlate with expanding cells, cell size should predict actin organization and vice versa. A root cell’s shape (length and width) or its actin array should correctly place the cell at a certain point in the expansion gradient of the elongation zone. While cell width could not predict any of the actin measurements—there was no predictive relationship between cell widths and actin filament density, skewness, angle, or parallelness (Figure 1C; Supplemental Figure 2)—cell lengths were highly predictive of each actin metric, using the descriptive statistical analysis bivariate fit (Figure 1D–G; Supplemental Figure 3). Short cells exhibited higher actin density (Figure 1D), lower bundling (Figure 1E), and what might be perceived as “disorganized” actin, with higher average filament angles (Figure 1F) and lower parallelness (Figure 1G) compared with long cells. These highly predictive relationships between cell length and each aspect of actin organization held when we examined actin organization in the Wassilewskija (WS) ecotype expressing GPF-fABD2 and in a T-DNA insertion mutant for the auxin transport protein AUX1 (*aux1-100*, WS background; Supplemental Figure 7), whose average root epidermal cell lengths are significantly longer than either wildtype. Although we were unable to accurately and consistently measure cell growth rates (based on literature such as Beemster and Baskin, 1998; van der Weele et al., 2003), and so have not determined whether bundles promote or even precede expansion, it is clear that a higher incidence of actin bundling occurred in long cells (Supplemental Figures 3, 7, and 9).

The parameter that adhered to a fairly linear relationship with cell length is parallelness (R^2^ = 0.68; Figure 1G; Supplemental Figure 3D)—how parallel filaments are to each other. Although these data cannot establish increased filament parallelness as the cause of cell elongation, they demonstrate that filament parallelness is the parameter most directly correlated with cell length. To determine whether any particular combination of the measured parameters (cell length, cell width, filament density, skewness, angle, and parallelness) explains the most variance from the mean for each cell, we performed principal component analysis on each data set, finding that the interactions between cell length, filament parallelness, and to a lesser extent, skewness, explain most of our observations for both wildtype ecotypes (Col-0 and WS) and the *aux1-100* mutant (see Supplemental Tables 1–6).

Aside from investigating correlations between actin organization and cell size, our intent was to find a more objective way of categorizing cells into “Region 2” or “Region 3” for wildtype plants. By plotting each cell’s specific actin metrics against its length or width, we defined maximum cell sizes for each region. The maximum length of a cell included as Region 2 became 85 µm, the mean cell length (57 µm) plus one standard deviation (28 µm); the minimum length of a cell included as Region 3 became 94 µm, the mean cell length (128 µm) minus one standard deviation (34 µm). These cutoffs were used in assigning “region” in all further experiments on the Col-0;GFP-fABD2 lines (see Methods).

### Cortical Actin Array Dynamics and Individual Filament Behaviors Differ between Short and Long Cells

Cortical actin arrays constantly remodel depending on the needs of a cell (Staiger et al., 2009; Henty et al., 2011; Henty-Ridilla et al., 2013; Henty-Ridilla et al., 2014; Cao et al., 2016). Arrays in isotropically-growing cotyledon pavement cells are observed to exhibit “more random” and “more dynamic” arrays than the anisotropically-growing cells of the root elongation zone (Smertenko et al., 2010). We hypothesized that the actin network in cells in Region 3 would be less dynamic than in Region 2. We collected 100-s timelapse movies from short and long cells in the same roots and calculated the pairwise correlation coefficient among all possible temporal intervals (Vidali et al., 2010). We found that the actin array dynamicity in Region 2 cells was significantly reduced compared to Region 3 (Supplemental Figure 4). The array of Region 2 cells was very dense, so we considered that a general comparison of pixel intensities and occupancies among temporal intervals of timelapse movies might not account for the true dynamic behavior of the array.

To determine what specific behaviors contribute to the overall filament array in cells, we quantified individual actin filament behaviors (Li et al., 2015b). We expected increased turnover in short cells and a higher frequency of bundling events in long cells. On average, filaments in short and long cells behaved similarly, except that longer cells exhibited longer, faster-growing filaments (Figure 2; Table 1). Upon measuring bundling, unbundling, and annealing frequencies, we were surprised to observe no differences in frequency of bundling or unbundling, but there was a multifold increase in annealing in shorter cells (Figure 2; Table 1).

**Table 1.**
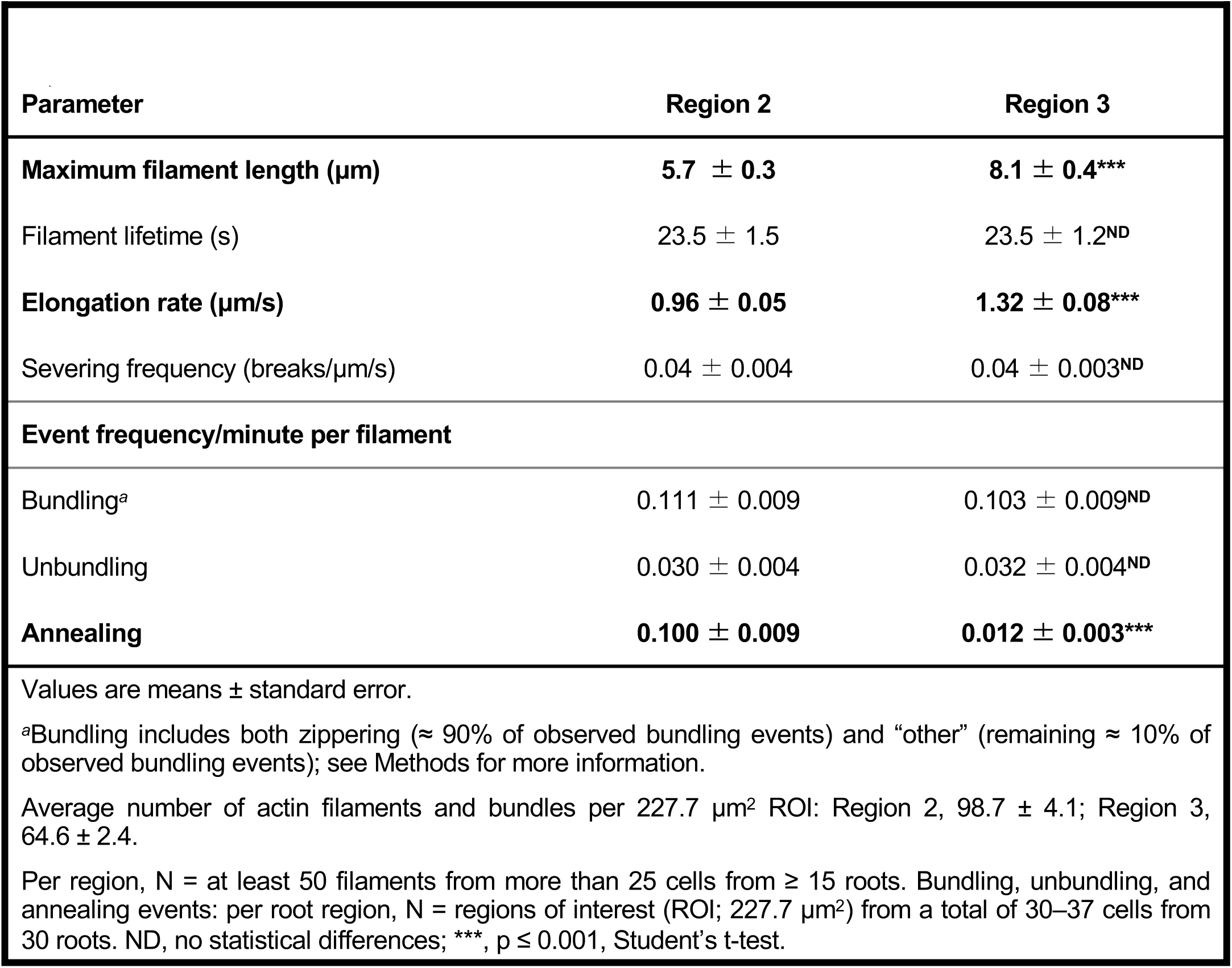
Individual Actin Filament Behaviors in Regions 2 and 3

| Parameter | Region 2 | Region 3 |
| --- | --- | --- |
| **Maximum filament length ($\mu\text{m}$)** | **$5.7 \pm 0.3$** | **$8.1 \pm 0.4^{***}$** |
| Filament lifetime (s) | $23.5 \pm 1.5$ | $23.5 \pm 1.2^{\text{ND}}$ |
| **Elongation rate ($\mu\text{m/s}$)** | **$0.96 \pm 0.05$** | **$1.32 \pm 0.08^{***}$** |
| Severing frequency (breaks/$\mu\text{m/s}$) | $0.04 \pm 0.004$ | $0.04 \pm 0.003^{\text{ND}}$ |
| **Event frequency/minute per filament** |  |  |
| Bundling<sup>a</sup> | $0.111 \pm 0.009$ | $0.103 \pm 0.009^{\text{ND}}$ |
| Unbundling | $0.030 \pm 0.004$ | $0.032 \pm 0.004^{\text{ND}}$ |
| **Annealing** | **$0.100 \pm 0.009$** | **$0.012 \pm 0.003^{***}$** |

Values are means ± standard error.
<sup>a</sup>Bundling includes both zippering ( $\approx 90\%$ of observed bundling events) and “other” (remaining $\approx 10\%$ of observed bundling events); see Methods for more information.
Average number of actin filaments and bundles per $227.7 \mu\text{m}^2$ ROI: Region 2, $98.7 \pm 4.1$ ; Region 3, $64.6 \pm 2.4$ .
Per region, N = at least 50 filaments from more than 25 cells from $\geq 15$ roots. Bundling, unbundling, and annealing events: per root region, N = regions of interest (ROI; $227.7 \mu\text{m}^2$ ) from a total of 30–37 cells from 30 roots. ND, no statistical differences; \*\*\*, $p \leq 0.001$ , Student’s t-test.

**Figure 2.**
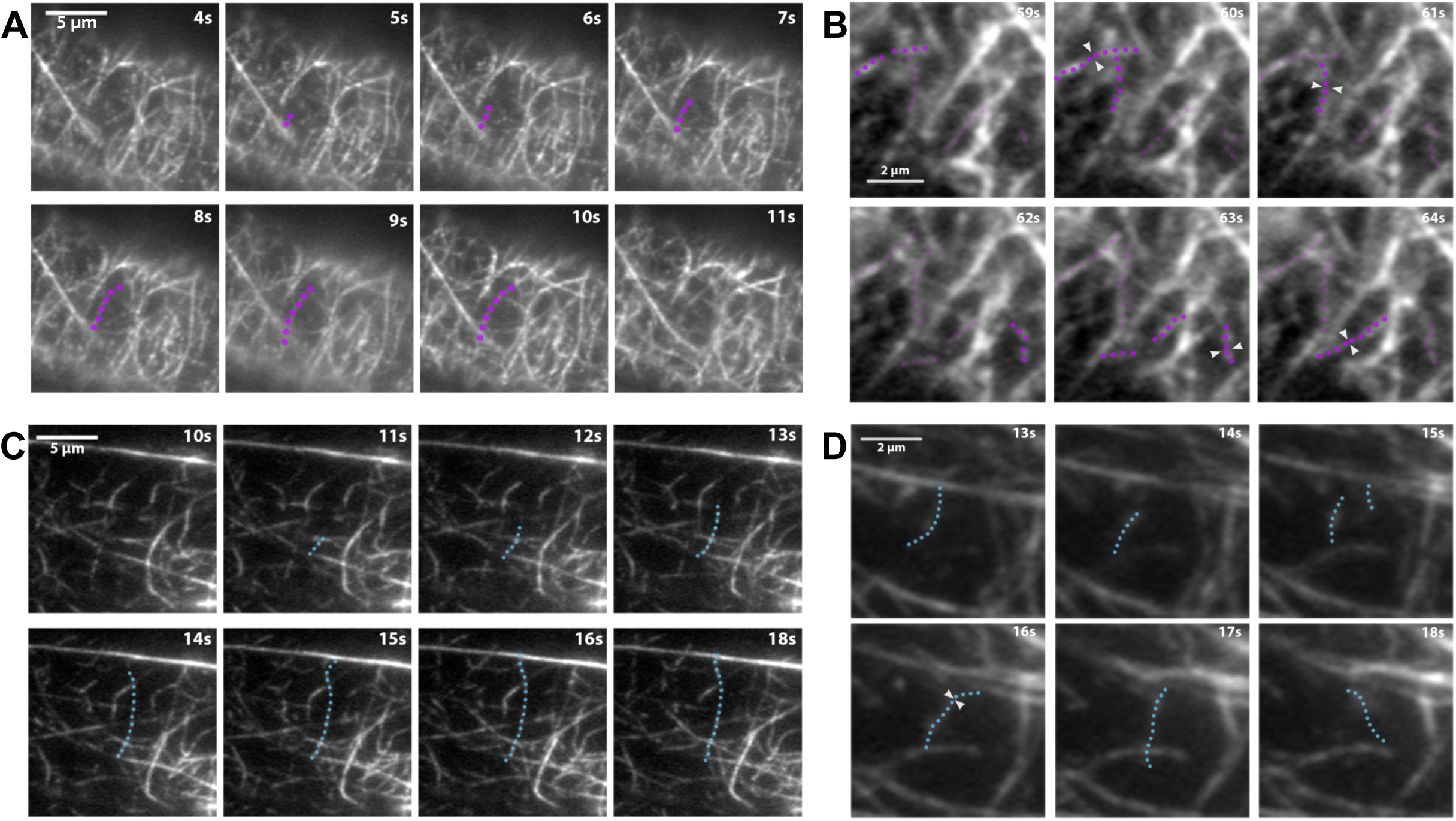
Timelapse Imaging of Cortical Actin Filaments in Root Epidermal Cells Shows Differences in the Dynamic Behavior between Short and Long Cells. **(A)** and **(C)** The cortical actin cytoskeleton in 6-day-old light-grown root epidermal cells expressing GFP-fABD2 was imaged with timelapse VAEM. Representative images of individual filament dynamics in short cells (up to 85 μm long, Region 2) and long cells (over 94 μm long, Region 3). On average, filaments in short cells **(A; filament highlighted in purple)** elongated over 25% more slowly and grew to be nearly 30% shorter than filaments in long cells **(C; filament highlighted in blue)**. Severing frequencies and filament lifetimes did not vary between regions; see Table 1. Scale bar, 5 μm. **(B)** and **(D)** Regions of interest (ROI; 227.7 μm) were selected from the same movies as **(A)** and **(C)**. Annealing occurs 10× more frequently in short cells **(B; filaments highlighted in purple)** compared with long cells **(D; filament highlighted in blue**). Note that four annealing events (white arrowheads) occurred within 6 s in **(B)** compared with only one event in **(D)**. Dots indicate fragments involved in annealing events. Quantification of annealing frequencies as well as bundling and unbundling frequencies are shown in **Table 1**. Although actin filament arrays in long cells were substantially more bundled compared with short cells (see Figure 1), there were no differences in bundling or unbundling frequencies when event frequencies were calculated on a per-minute, per-filament basis. Scale bar, 2 μm. 100-s timelapse movies were collected from short and long cells in the same 30 roots. Note: Brightness and contrast were enhanced in the montages of Figure 2B and Figure 2C to better show the filament and its changes.

### Actin Organization Responds to Short-Term IAA Treatments

To decipher which actin parameter(s) coincided with cell expansion, and to find stronger indicators of causality, we treated roots with a known inhibitor of root growth, the naturally occurring auxin, IAA, which has been shown to inhibit root growth within minutes of application (Hejnowicz and Erickson, 1968; Fendrych et al., 2018). Auxin affects actin organization and dynamics and inhibits root growth (reviewed in Zhu and Geisler, 2015; Fendrych et al., 2018), known to depend on an intact cytoskeleton (reviewed in Hussey, 2006 and Li et al., 2015a). Yet, actin response following short-term auxin treatments, i.e., a way to directly link the two, has not been determined. If decreased actin density and increased bundling are indeed hallmarks of growth, and if auxin works by modulating the actin cytoskeleton, then IAA, an agent that inhibits growth, should induce the opposite actin phenotype: after IAA treatment, density should increase and bundling decrease. Further, if a lower average filament angle and higher parallelness are indicative of rapidly growing cells (described in Dyachok at al., 2011, and a natural assumption given that we found increasingly longitudinal actin arrays in longer cells), applying IAA should increase filament angle and decrease parallelness (i.e., there should be a decrease in both longitudinality and apparent “organization”).

As expected, 20–30 min IAA treatments induced significant increases in actin filament density and decreases in extent of bundling (Figure 3), linking actin abundance and a reduction in bundling with plant response to IAA. We were surprised, however, to observe a dose-dependent increase in apparent actin organization after IAA treatments (Figure 3A,D–E, Supplemental Figure 5). In another, time series experiment, we established that the IAA-induced increase in parallel longitudinality is maintained for at least 60 min after initial treatment (Supplemental Figure 6). Strong changes in filament angle and orientation generally appeared more slowly than the increase in density. Together, these are the first data that quantitatively document actin’s short-term response to moderate doses of IAA.

**Figure 3.**
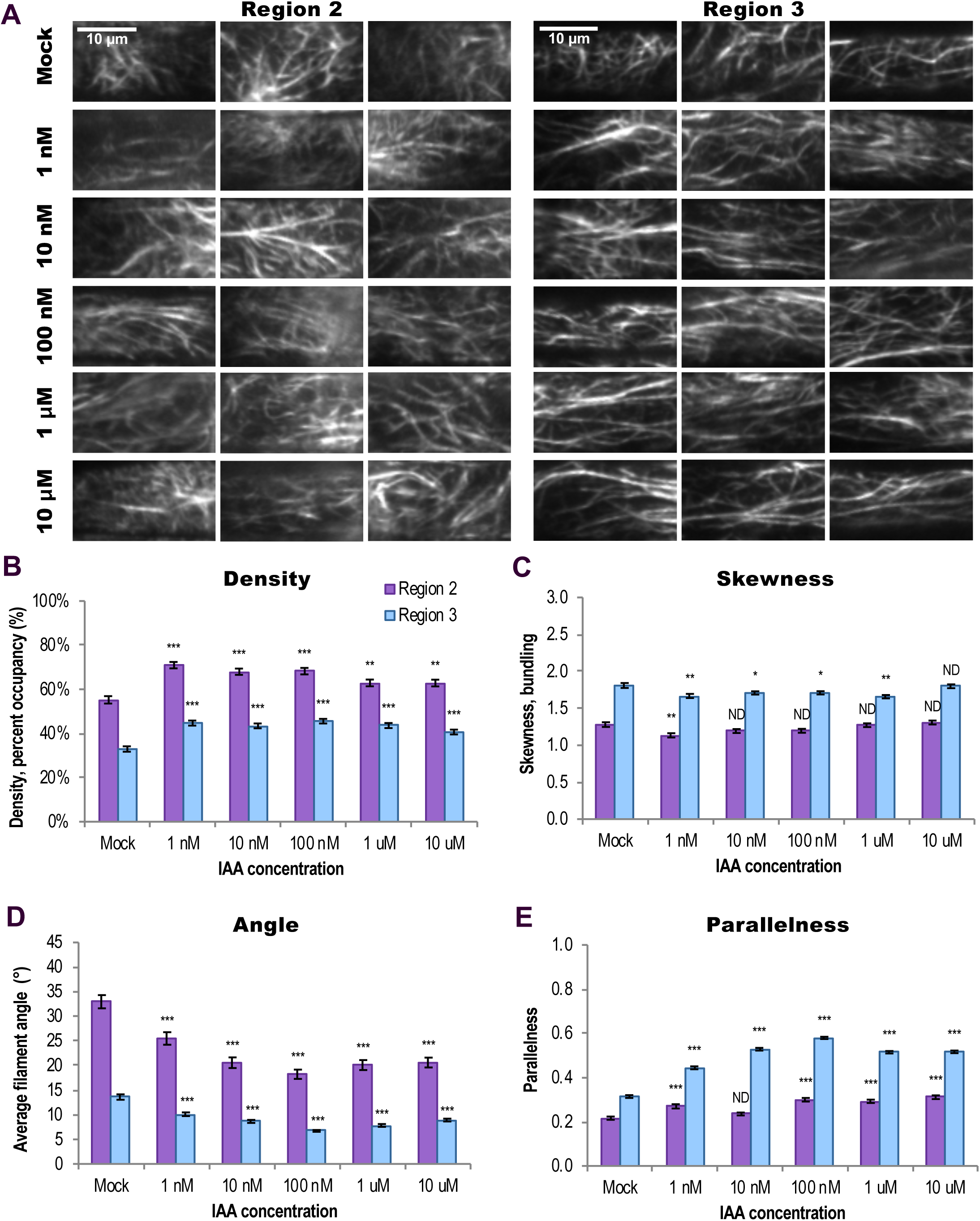
Short-Term IAA treatments Induce Changes in Actin Filament Organization. **(A)** Representative VAEM images of GFP-fABD2–labeled actin in epidermal cells from Region 2 (≤ 85 μm long) and Region 3 (≥ 94 μm long), treated for 20–30 min with indicated doses of IAA or mock. Scale bar, 10 μm. **(B)** to **(E)** Quantification of actin architecture and orientation in root epidermal cells: IAA triggered an increase in actin filament density **(B)** and decrease in skewness **(C)**. Region 2 measurements are shown in purple; Region 3 in blue. **(D) and (E)** After IAA treatments, actin arrays in both regions were more “organized,” with lower average filament angle **(D)** relative to the longitudinal axis of the cell and filaments generally more parallel to each other **(E)**. Changes in actin orientation **(D)** and **(E)** were dose-dependent, see Supplemental Figure 5. Cells whose lengths fell between 85 and 94 μm were counted in both regions. N = 8–12 cells per region per root from at least 10 roots per treatment. ND, no statistical differences; *, p ≤ 0.05; **, p ≤ 0.01; ***, p ≤ 0.0001, oneway ANOVA, compared with Dunnett’s Method, comparing doses to mock in each Region, in JMP. Results are from one representative experiment of 3 similar experiments with similar results. All IAA experiments were performed and analyzed double blind.

### Actin Filaments Unbundle in Response to IAA

Links between auxin and actin clearly exist (reviewed in Zhu and Geisler, 2015) but the specific components of these pathways—and actin’s role in them—are unresolved, so we evaluated actin’s role in IAA perception by measuring whether individual filament behaviors change in the minutes immediately following IAA treatment.

Since substantial increases in actin density and parallel longitudinality occurred within 20–30 min of treatment with IAA, we hypothesized that individual filaments would respond quickly to treatment and might undergo increased severing, faster filament elongation rates, increased unbundling, and/or increased annealing (a result of decreased end-capping; Li et al., 2012). We tested this prediction by quantifying actin dynamics in epidermal cells in Region 2 and Region 3 within 7 min of 10 nM IAA treatment. Surprisingly, we observed no changes in most individual filament behaviors in either shorter or longer cells within this 7-min timeframe (Table 2). Though the differences in filament elongation rates and maximum filament length we previously observed between regions (Table 1) were reproduced, 10 nM IAA did not affect any of the measured stochastic dynamics parameters: overall filament length, lifetime, elongation rate, or severing frequency within a region. However, when we measured frequency of bundling, unbundling, and annealing, we observed an IAA-induced doubling of unbundling events in both short and long cells (Figure 4, Table 2). In long cells, IAA induced a near 5-fold increase in annealing (Figure 4D, Table 2). Actin filaments unbundled and altered annealing frequencies within 7 min of IAA treatment, demonstrating that actin participates in short-term responses to the hormone.

**Figure 4.**
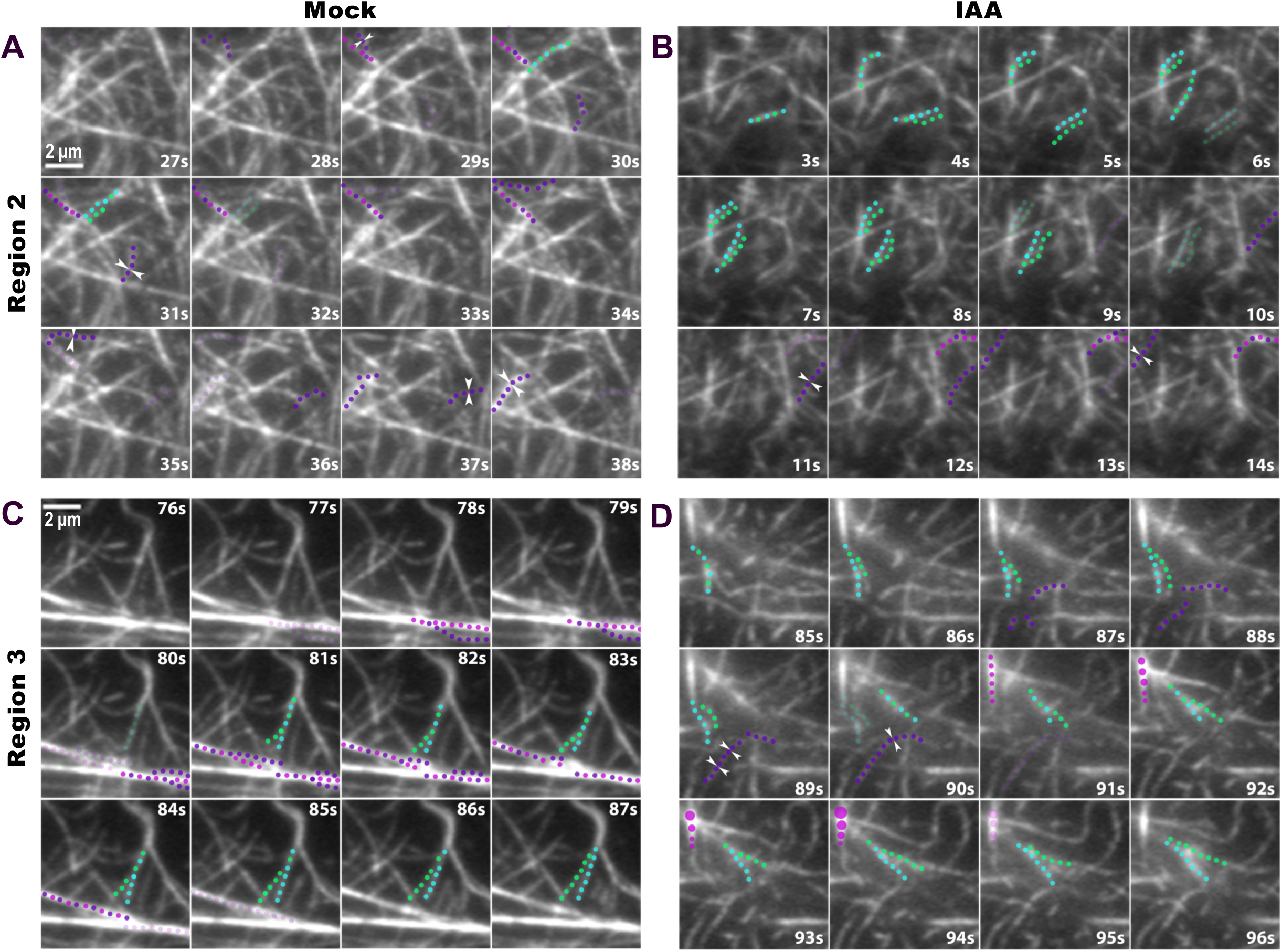
Short-Term Auxin Treatments Cause Actin Filament Unbundling. Representative images of individual filament bundling, unbundling, and annealing in Region 2 cells **(A)** and **(B)** and Region 3 cells **(C)** and **(D)**; mock **(A)** and **(C)** vs. 10 nM IAA **(B)** and **(D)**. Scale bar, 2 μm. **(A)** and **(B)** Timelapse series of VAEM images show that 10 nM IAA **(B)** increased actin filament unbundling in Region 2 within 7 min compared with mock **(A)**. Note that one unbundling event (filament unbundling shown as blue and green dots separating) occurred in **(A)** whereas three occurred in the same timespan in **(B)**. There was also a small but statistically significant decrease in annealing events (white arrowheads) after IAA treatment. Other aspects of individual filament behaviors did not significantly change after treatment; for complete quantification of all measured individual filament dynamics, see Table 2. **(C)** and **(D)** Treatment with 10 nM IAA **(D)** increased actin filament unbundling and filament end annealing in Region 3 within 7 min compared with mock **(C)**. Similarly as in Region 2, IAA stimulated unbundling of actin filaments: two unbundling events are shown in **(D)** compared with only one event in **(C)**. IAA also stimulated an increase in annealing in Region 3, where three annealing events are shown by white arrowheads **(D)**. Bundling events are shown by either purple and magenta dots coming together (zippering of two independent filaments) or a series of magenta dots increasing in size (fluorescence intensity increase with no visible filament zippering). 100-s timelapse movies were collected from short and long cells in the same 28 6-day-old, light-grown roots.

**Table 2.** Actin Filament Dynamics after Treatment with IAA

|  | Region 2 |  | Region 3 |  |
| --- | --- | --- | --- | --- |
| Parameter/Treatment | MOCK | IAA | MOCK | IAA |
| Maximum filament length ($\mu\text{m}$) | $5.2 \pm 0.2$ | $4.6 \pm 0.2^{\text{ND}}$ | $9.8 \pm 0.6$ | $9.7 \pm 0.4^{\text{ND}}$ |
| Filament lifetime (s) | $27.4 \pm 1.9$ | $24.9 \pm 1.2^{\text{ND}}$ | $28.8 \pm 2.2$ | $33.9 \pm 2.1^{\text{ND}}$ |
| Elongation rate ($\mu\text{m/s}$) | $0.97 \pm 0.14$ | $0.88 \pm 0.05^{\text{ND}}$ | $1.63 \pm 0.06$ | $1.62 \pm 0.07^{\text{ND}}$ |
| Severing frequency (breaks/$\mu\text{m/s}$) | $0.05 \pm 0.003$ | $0.06 \pm 0.003^{\text{ND}}$ | $0.03 \pm 0.002$ | $0.03 \pm 0.002^{\text{ND}}$ |
| **Event frequency/minute per filament** |  |  |  |  |
| Bundling<sup>a</sup> | $0.216 \pm 0.023$ | $0.170 \pm 0.015^{\text{ND}}$ | $0.239 \pm 0.023$ | $0.187 \pm 0.015^{\text{ND}}$ |
| Unbundling | **$0.092 \pm 0.010$** | **$0.218 \pm 0.017^{***}$** | **$0.104 \pm 0.012$** | **$0.208 \pm 0.025^{**}$** |
| Annealing | **$0.172 \pm 0.009$** | **$0.133 \pm 0.013^*$** | **$0.034 \pm 0.006$** | **$0.147 \pm 0.020^{***}$** |
| Values are means ± standard error. | | | | |
| <sup>a</sup>Bundling includes both zippering ($\approx 90\%$ of observed bundling events) and “other” (remaining $\approx 10\%$ of observed bundling events); see Methods for more information. These percentages hold for both mock- and IAA-treated plants. | | | | |
| Average number of actin filaments and bundles per $57.8 \mu\text{m}^2$ ROI: Region 2, $48.6 \pm 2.1$; Region 2 + IAA, $48.4 \pm 1.6$; Region 3, $26.9 \pm 1.1$; Region 3 + IAA, $28 \pm 1.3$; see Methods. | | | | |
| 6-day-old roots were treated with 10 nM IAA or mock and epidermal cells were imaged for up to 7 min after treatment. |  |  |  |  |
| Per region per treatment, N = at least 50 filaments from $\geq 20$ cells from $\geq 12$ roots. ND = no statistical differences, Student's t-test compared with mock for that region. Bundling, unbundling, and annealing events: per root region per treatment, N = regions of interest (ROI; $57.8 \mu\text{m}^2$) from a total of 21–23 cells from 18–22 roots. *, $p \leq 0.05$; **, $p \leq 0.001$; ***, $p \leq 0.0001$; ND, no statistical differences, Student's t-test. | | | | |

### The Actin Array in *aux1* Mutants is Insensitive to IAA but Partially Responds to NAA

The auxin importer AUX1 was identified in an ethyl methanesulphonate (EMS) mutant screen for resistance to IAA and 2,4-D (Maher and Martindale, 1980; Pickett et al., 1990), and the protein has been shown to bind IAA with extremely high affinity (Yang et al., 2006; Carrier et al., 2008). AUX1 mutants are agravitropic and exhibit root elongation in the presence of the natural auxin IAA (whereas wildtype roots are growth-inhibited under this condition), but mutant root growth is inhibited to wildtype levels in the presence of the membrane permeable auxin NAA (Maher and Martindale, 1980; Pickett et al., 1990; Bennett et al., 1996; Marchant et al., 1999). Cells in *aux1* mutants take up significantly less IAA (Rashotte et al., 2003; Hayashi et al., 2014; Rutschow et al., 2014, protoplasts; Dindas et al., 2018) and are larger compared with wildtype cells (Ugartechea-Chirino et al., 2010; Supplemental Figures 7 and 9). Intracellular auxin concentrations correlate with cell length, where IAA concentrations are higher in longer root epidermal cells (Brunoud et al., 2012), possibly because IAA concentration regulates the amount of time cells spend in the elongation zone (Rahman et al., 2007). We hypothesized that AUX1 might have a previously uncharacterized role in short-term auxin signaling to the cytoskeleton.

We expressed GFP-fABD2 in the T-DNA insertion mutant for AUX1, *aux1*-*100* (WS background) and in the point mutant *aux1*-*22* (Col-0 background; Feldmann, 1991; Roman et al., 1995; Bennett et al., 1996), with the hypothesis that if AUX1 were upstream of cytoskeletal rearrangements in response to IAA, the mutants’ actin cytoskeleton would not respond to 20–30 min IAA treatments: neither density nor parallelness would increase, and neither skewness nor average filament angle would decrease. Because root epidermal cells in *aux1* plants were significantly longer than wildtype (Supplemental Figures 7 and 9), analyzing actin response by separating cells into standard “regions” seemed imprecise. For example, a 120 μm-long cell that will grow to a final length of 140 μm in a wildtype plant is at a different point in its development than a 120 μm-long *aux1*-*100* cell that will reach a final length of 290 μm. Therefore, we quantified changes in actin array on a per cell basis (see Methods).

Both *aux1* mutants had average cell lengths longer than wildtype, as well as overall actin array organization that differed from wildtype under control conditions (Figure 5 and Supplemental Figures 7, 8, and 9). When each mutant was compared to its respective wildtype ecotype, both alleles of *aux1* exhibited significantly lower average filament density and increased skewness/bundling. Filaments were overall more longitudinal and parallel to one another. Mutants’ longer cells and “more organized” actin filament organization fits the model that higher levels of apparent “organization” are coincident to cell expansion.

**Figure 5.**
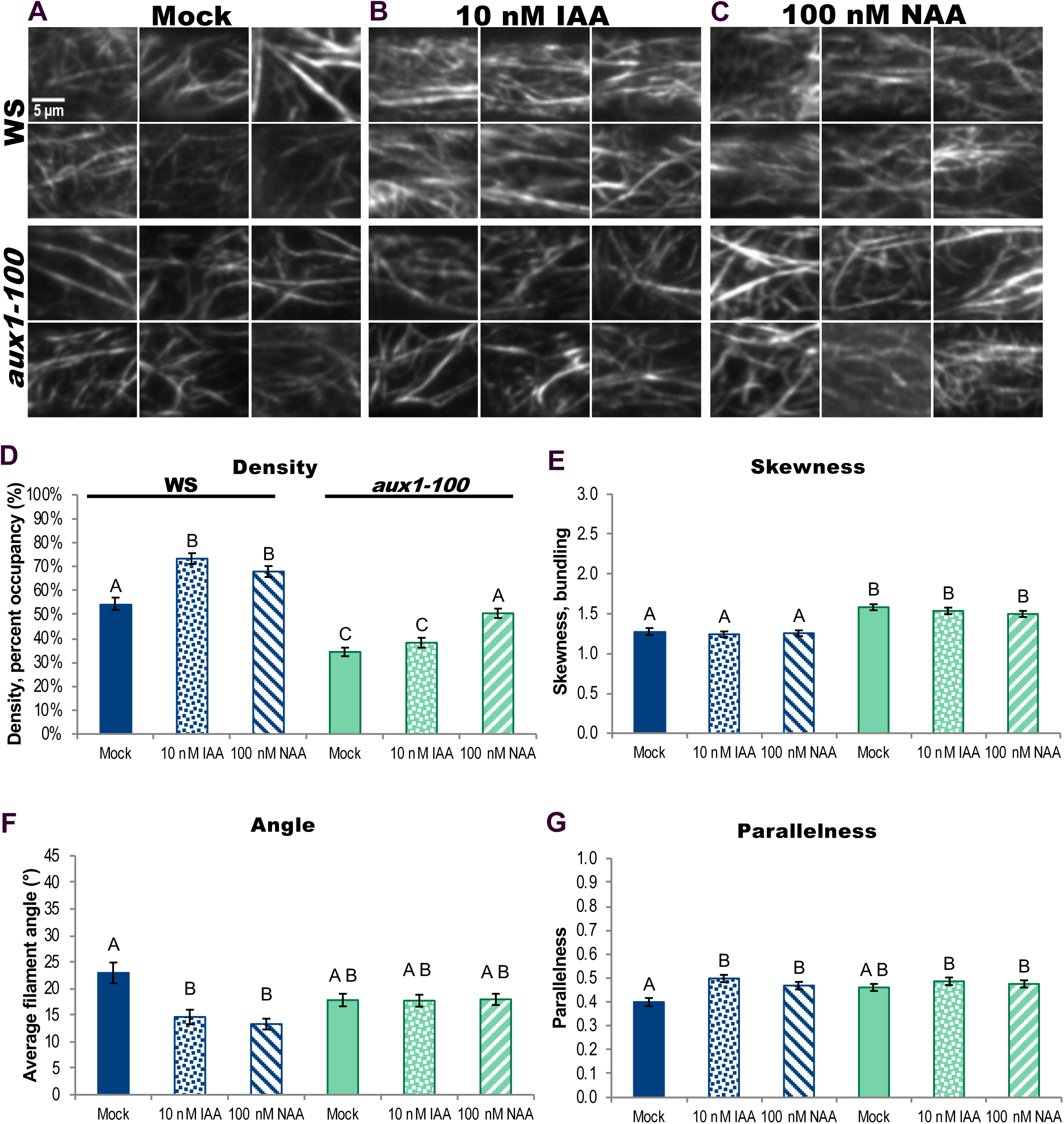
Actin Organization in *aux1-100* Fails to Respond to Short-Term IAA Treatments but Partially Responds to the Membrane-Permeable Auxin NAA. **(A)** to **(C)** Representative VAEM images of GFP-fABD2–labeled actin in epidermal cells from wildtype and *aux1*-*100*, treated for 20–30 min with mock **(A)**, 10 nM IAA **(B)**, or 100 nM NAA **(C)**. Scale bar, 5 μm. **(D)** to **(G)** Quantification of actin organization in root epidermal cells. IAA failed to trigger an increase in actin filament density in *aux1*-*100* **(D)** but actin density in *aux1-100* increased in response to NAA. Skewness in both genotypes did not significantly respond to either treatment **(E)**. Wildtype response is shown in blue and *aux1*-*100* in green; mock, solid; 10 nM IAA, dots; 100 nM NAA, stripes. After IAA and NAA treatments, actin arrays in wildtype plants were more “organized,” with lower average filament angle **(F)** relative to the longitudinal axis of the cell and filaments generally more parallel to each other **(G)**. Average actin filament angle and parallelness in *aux1*-*100* failed to reorganize in response to either IAA (dots) or the membrane-permeable auxin NAA (stripes). N = 7–32 cells per root; 9–11 roots per genotype per treatment. Different letters indicate statistically significant differences, oneway ANOVA, compared with Tukey-Kramer HSD in JMP. Actin measurements were quantified on a per-cell basis; see Methods for description and Supplemental Figure 7 for scatter plots. Results are from one representative experiment of 2 similar experiments with similar results. All auxin experiments were performed and analyzed double blind.

Actin organization in wildtype WS plants expressing GFP-fABD2 responded to short-term IAA treatments almost identically as had Col-0. Actin filament density significantly increased, as did parallelness, and average filament angle significantly decreased (Figure 5). Interestingly, when actin response was quantified on a per cell basis, WS did not exhibit the small but statistically significant decrease in skewness/bundling (Figure 5E and Supplemental Figure 7) that we had previously observed in Col-0 (Figure 3), perhaps because the ecotype itself is slightly resistant to auxin (Dharmasiri et al., 2005b). The a*ux1*-*100* mutant’s actin array did not significantly reorganize in response to IAA treatment (Figure 5), indicating that actin cytoskeleton response to IAA required the transporter. To confirm the importance of AUX1 in IAA-triggered actin cytoskeleton rearrangements, we tested a second allele, the null point mutant *aux1*-*22*. The actin array in *aux1*-*22* also failed to reorganize in response to IAA treatments (Supplemental Figure 8).

To understand whether the auxin hormone itself drives cytoskeletal reorganization, or if there is an intermediary between auxin, AUX1, and actin response, we tested the mutant’s response to the membrane permeable auxin NAA. If AUX1’s role is restricted to transporting IAA into the cell and auxin itself merely needs to enter the cell to stimulate actin reorganization, NAA should be sufficient to induce a wildtype response in *aux1*-*100* and we should see denser, more parallel, and more longitudinal arrays. But if NAA should fail to induce the established reorganization pattern, we could deduce that the presence of auxin inside the cell is not enough and that the AUX1 protein is required for short-term auxin to actin signaling. Figure 5 shows that WS responds to NAA similarly as to IAA. Interestingly, NAA only partially restores in *aux1-100* a wildtype response to IAA. NAA stimulates increased actin filament density (Figure 5D) in the mutant, but has no effect on filament angle or parallelness (Figures 5F–G).

To confirm AUX1’s importance in cytoskeletal responses to NAA, we tested the membrane permeable hormone’s effects on actin organization in *aux1-22* and its wildtype, Col-0. Recapitulating *aux1-22*’s lack of response to IAA, actin organization in this mutant was largely impervious to NAA, with a sizeable but statistically insignificant reduction in average filament angle. Upon testing the effect of NAA on actin reorganization in Col-0, we found that NAA stimulated an increase in actin filament density in Col-0, but were surprised that, when measured on a per cell basis, Col-0 exhibited neither a decrease in average filament angle nor an increase in filament parallelness (Supplemental Figure 8). The Col-0 and WS ecotypes are likely genetically divergent enough to explain why their NAA-prompted actin reorganization is not identical; indeed, previous work identified transcriptional responses for several genes that differ among the two ecotypes under various environmental conditions, including for other proteins involved in hormone signaling (Schultz et al., 2017). Actin organization in both *aux1-100* and *aux1-22* failed to respond to IAA and only partially responded to NAA. These results are the first that place AUX1 upstream of actin in short-term auxin signaling events, as well as demonstrate that the import protein is required for a complete actin response to auxin.

## DISCUSSION

We correlated specific actin architecture and orientation to cell lengths in expanding root epidermal cells and report the first quantitative assessment of actin responses to short-term IAA treatments in roots. Under control conditions, short epidermal cells (Region 2) are characterized by dense actin arrays with high annealing frequencies, whereas long cells (Region 3) exhibit more bundled, more parallel, more longitudinal actin arrays in which filaments elongate faster and grow longer. We found that this same pattern of actin organization occurs in the WS ecotype and *aux1* mutants, indicating that there may be a causal relationship among cell length, skewness/bundling, and filament parallelness. We documented actin responses to growth-inhibitory doses of IAA and were surprised to find that filaments became more dense, parallel, and longitudinally oriented (i.e., lower average filament angle) within 20–30 min, demonstrating that the relationship between higher levels of actin “organization” and increased cell expansion is not as direct as previously hypothesized. Upon analyzing the actin array response to auxin in two *aux1* mutants (the T-DNA insertion mutant *aux1-100* and the null point mutant *aux1-22*), we found that actin failed to reorganize in response to IAA and actin reorganization was only partially restored by NAA. Although none of our results establishes a cause-and-effect relationship between increased actin bundling and elongating cells, they disprove the hypothesis that actin bundles inherently inhibit cell expansion. Although some specific actin characteristics correlate with longer cells, “more organized” filament arrays do not universally correlate with rapidly growing root cells. We also provide the first evidence that the auxin import protein AUX1 is critical for the actin cytoskeleton’s full response to short-term auxin treatments, presumably because it imports the bulk of IAA into cells, and that cytoplasmic auxin is sufficient to trigger some aspects of actin reorganization.

Studies on the long-term (6+ hours) effects of high doses of auxin on actin filaments (Li et al., 2014a; Scheuring et al., 2016) report that actin becomes more bundled after treatment. Several studies (Holweg et al., 2004; Nick et al., 2009) examining the effect of short-term, high dose IAA treatments on m-Talin–bundled actin filaments in dark-grown rice coleoptiles demonstrate that relatively high doses of auxin (10 µM NAA, or 50 µM IAA) induced filament unbundling. Auxin transport inhibitors appear to have the opposite effect on actin filaments. Auxin transport inhibitors such as 2,3,5-triiodobenzoic acid (TIBA) induce actin bundling within minutes (Dhonukshe et al., 2008) and inhibit cell elongation (Rahman et al., 2007). Here, we demonstrated that short-term treatment with IAA, closer to endogenous levels (Band et al., 2012), induced an increase in filament density, parallelness, and longitudinality, and decrease in bundles within 20–30 min, as well as an increase in unbundling events in cells throughout the visible root elongation zone within 7 min (Table 2). These results indicate that, like their role in microbe-associated molecular pattern (MAMP) perception (Cárdenas et al., 1998; Henty-Ridilla et al., 2014; Li et al., 2015b), actin filaments are potentially involved in the initial intracellular perception of IAA. Interestingly, unbundling actin filaments to build a dense array in auxin signaling/response is a different cellular mechanism used to increase filament density vs. the increased density observed in responses to MAMPs in Arabidopsis hypocotyls, where MAMPs inhibit actin depolymerizing factor (ADF)-mediated severing and downregulate capping protein-(CP) mediated barbed end capping to build a denser array (Henty-Ridilla et al., 2014; Li et al., 2014b). This indicates that, despite similar actin readouts of “increased density” after MAMP or 20 min IAA treatment, actin participates in discrete roles in each of these signaling–response pathways, and each stimulus distinctly modulates actin regulation towards separate, precise outcomes.

We found that the auxin import protein AUX1 is required for short-term changes in actin organization in response to auxin. Previously the role of “auxin receptor” was attributed to ABP1 (Chen et al., 2001; Chen et al., 2012; Lin et al., 2012; Nagawa et al., 2012; reviewed in Sauer and Kleine-Vehn, 2011), but immediate cytoskeletal reorganization was never directly linked to the protein because though actin response in roots was implied (Lin et al., 2012), it was never directly visualized. The *aux1* mutants’ actin arrays are impervious to IAA, likely because auxin cannot enter cells in sufficient quantities, and are only partially responsive to the membrane-permeable NAA, indicating that while auxin itself elicits some actin rearrangements, AUX1 is necessary for full response. Having an established baseline of auxin’s short-term effects on actin filaments in wild-type and *aux1* root epidermal cells will enable further testing of models of actin’s role in auxin signaling pathways.

### Actin Organization Predicts Cell Length Under Control Conditions but Not Otherwise

Actin’s role in cell expansion is established but not understood (reviewed in Li et al., 2015a). It has been generally accepted that cell expansion requires a degree of observable actin “organization” (Smertenko et al., 2010; Dyachok et al., 2011). The hypothesis that an “organized” actin array corresponds with cell growth appeals to an intuitive understanding of growth as a methodical process that requires coordinated elements. However, “organization” can take many forms and exactly what form drives growth is unknown. Indeed, we found in two ecotypes—and in an auxin signaling mutant with significantly longer cells—that filament parallelness, cell length, and skewness affect one another to a similar extent to produce predictable organization across the root elongation zone. By showing that, under control conditions, actin bundling increases as cell length increases, we present definitive evidence that filament bundles do not inhibit cell expansion. Our observation, substantiated by quantitative evidence, that the cytoskeleton changes to a higher level of apparent actin “organization”, namely longitudinality and filament parallelness, in response to IAA, a treatment known to inhibit growth within minutes (Hejnowicz and Erickson, 1968; Fendrych et al., 2018), definitively demonstrates that increases in filament “organization” do not, inherently, contribute to cell expansion. Furthermore, the IAA-induced decrease in overall filament bundling (Figure 3c) and increase in unbundling events (Table 2) that occurs in Col-0 indicates that an absence of longitudinal bundling does not necessarily coincide with expanding cells. Thus, although our data cannot divulge a cause-and-effect relationship, our correlative study has eliminated two hypotheses for the relationship between actin organization and cell expansion and determined that there is no absolute relationship between actin bundling and cell expansion.

### Actin Behaviors Differ in Short and Long Epidermal Cells of the Root Elongation Zone

We evaluated individual filament behaviors in short and long cells to gain insight into what filament behaviors might contribute to cell growth. Although several aspects of individual filament dynamics were previously measured in root epidermal cells in Arabidopsis (Smertenko et al, 2010), we have examined additional filament behaviors, bringing knowledge of Arabidopsis up to that of rice (Wu et al., 2015). Region 2 was significantly denser than Region 3 so we expected a higher rate of filament turnover: increased severing, shorter filaments and filament lifetimes, and faster elongation rates. Filament arrays in Region 3’s longer cells were much more bundled than arrays in Region 2 so it was reasonable to expect either a higher bundling frequency in longer cells, a higher incidence of unbundling in shorter cells, or a lower incidence of unbundling in longer cells. In this study, both short and long cells exhibited similar individual filament behaviors (Table 1), the only major differences being a reduced maximum filament length and elongation rate in shorter cells, and a multifold increase in incidents of annealing in shorter cells. In etiolated hypocotyls, increased filament lengths and lifetimes correlate with longer cells (Henty-Ridilla et al., 2013; Li et al., 2014b). Although shorter root epidermal cells have shorter average filament lengths, lifetime is statistically equivalent to that of filaments in longer cells, demonstrating that the connection between longer lifetime and increased cell length is not evident in roots.

The most marked difference in individual filament behaviors between short cells and long cells was the up to–10-fold increase in annealing frequency observed in shorter cells. Although the correlation coefficient algorithm found Region 2 to be less dynamic than Region 3, pixels occupied by fluorescence do not change as much during annealing events as, for example, when an entirely new filament polymerizes, so it is possible that this method failed to capture the full range of dynamic behaviors in Region 2 cells’ actin arrays. The substantial increase in annealing on a per-filament basis indicates there is likely a purpose behind this phenomenon. Annealing in vitro is generally a function of actin concentration, filament length, and/or filament end availability (Adrianantoandro et al., 2001), and in vivo is down-regulated by capping protein (Henty-Ridilla et al., 2014; Li et al., 2014b). Annealing is a way to build filaments quickly and without intensive energy inputs (Smertenko et al., 2010; Li et al., 2014b). Since maximum filament length is reduced in shorter cells that have a higher annealing frequency compared with maximum filament length in longer cells, and since these seem to be transient annealing events that for the most part hold for only a few frames, the purpose of these events does not appear to be building longer filaments. Perhaps these shorter cells, which are located in the subsection of the elongation zone known as the transition zone, are undergoing more cytoplasmic changes in preparation for rapid elongation.

### AUX1 is Necessary for Actin Response to Auxin

It is well-established that signaling between auxin and actin occurs (Kleine-Vehn and Friml, 2008; Nick et al., 2009; Nick, 2010; Lin et al., 2012; Nagawa et al., 2012; Li et al, 2015a; Scheuring et al., 2016; Zhu et al., 2016), but how auxin affects growth, how auxin affects actin, and how auxin affects actin to influence cell expansion are far from being understood. The multiple auxin–actin pathways (reviewed in Overvoode et al., 2010 and Grones and Friml, 2015) assign various roles to actin in auxin response (for example, repositioning auxin transport proteins, or inhibiting endocytosis of auxin transporters), and the pathway thought to link auxin and actin in the very short-term was via the plasma membrane auxin receptor ABP1 (Xu et al., 2010, 2011; Nagawa et al., 2012). Now that ABP1’s role in auxin–actin signaling is in doubt (Dai et al., 2015; Gao et al., 2015), the mechanism of auxin–actin signaling and how actin rapidly perceives auxin—through an upstream receptor, second messenger(s) modulating actin binding proteins, and/or perhaps direct interaction with the hormone itself—remains undetermined.

Before this current work, AUX1 was not suspected of playing a “transceptor” role in signaling upstream of actin organization. AUX1 is an established transporter of auxin (Bennett et al., 1996; Dindas et al., 2018), has homologs in all plants (reviewed in Swarup and Péret, 2012), is necessary for root gravitropism (Maher and Martindale, 1980; Marchant et al., 1999; Swarup et al., 2004), and is responsible for 80% of IAA uptake in root hairs (Dindas et al., 2018) and rapid growth inhibition by IAA (Fendrych et al., 2018). Auxin-induced transcriptional regulation requires that auxin binds to the intracellular auxin receptors SCF^TIR1/AFB^, which complex then binds to AUXIN/INDOLE-3-ACETIC ACID (Aux/IAA) transcriptional repressors (Dharmasiri et al., 2005a,b; Calderón Villalobos et al., 2012). However, the SCF^TIR1/AFB^ pathway is also responsible for short-term intracellular responses to auxin. Within the first 10 min of receiving an auxin signal, there is both an initial influx of H^+^ (depolarizing the plasma membrane and reducing cytosolic pH) and, shortly thereafter, increased intracellular Ca^2+^ that propagates through the root (Dindas et al., 2018). Both IAA and NAA are substrates of AUX1 (Yang et al., 2006; Carrier et al., 2008), and of the SCF^TIR1/AFB^–Aux/IAA complex, though both AUX1 and the protein complex have a higher affinity for IAA (Dharmasiri et al., 2005b; Calderón-Villalobos et al., 2012; Dindas et al., 2018).

Actin in wildtype cells reorganizes to increase filament density in response to both IAA and NAA. Surprisingly, NAA stimulated different effects on actin reorganization in Col-0 and WS: WS responded to NAA as it had to IAA but Col-0 underwent only increased actin filament density, and no changes in parallel longitudinality at 20–30 min. Col-0 might respond to NAA on a timeframe different from WS and different from its response to IAA. Natural and synthetic auxins inhibit root elongation to different extents, depending on ecotype (Delker et al., 2010). Occasionally differential responses to NAA have been detected across ecotypes, for example NAA stimulates a higher number of lateral roots in 9-day-old Col-0 plants compared with WS or Landsberg erecta (Falasca and Altamura, 2003). In addition, IAA and NAA induce in Col-0 different extents of gene expression (Yoshimitsu et al., 2011). It is quite possible that the two ecotypes are more different than is commonly acknowledged, a subject that merits further study.

Actin reorganization in both alleles of *aux1* was resistant to IAA and exhibited only partial responses to NAA, implicating AUX1 as a major player in auxin signaling to actin. After NAA treatments, density increased only in *aux1-100* and angle decreased (although not statistically significantly) only in *aux1-22*. Our results that show attenuated actin reorganization in *aux1-100’s* and *aux1-22*’s responses to NAA support the Dindas et al. (2018) model of an intracellular feedback loop that relies on the presence of AUX1. In the *aux1* mutant *wav5-33*, NAA triggers membrane depolarization (at a level similar to wildtype) and H^+^ influx (though substantially delayed and reduced vs. wildtype), but all *aux1* mutants tested are severely or entirely resistant to IAA in these responses (Dindas et al., 2018). Nor does IAA treatment in *aux1* mutants lead to the typical increased Ca^2+^ and although NAA was not directly tested (Dindas et al., 2018), these results imply that NAA might not induce Ca^2+^ influx in *aux1* mutants. Both H^+^, which contributes to membrane depolarization, and Ca^2+^ are known regulators of various actin binding proteins (see below). It is possible that import of IAA through AUX1 is necessary to activate or inhibit other intracellular players which drives, separately, increased actin density, decreased filament angle, and increased parallelness. Alternatively, actin reorganization might be delayed in the mutants. In any case, that previous study (Dindas et al., 2018) and our data showing partial actin response to NAA in *aux1* mutants demonstrate that auxin itself can act as an intracellular signaling molecule that stimulates some short-term cellular responses, a deviation from the ABP1 model that relied on IAA being perceived at the plasma membrane and its signal amplified within the cell.

### Potential Players in the Actin–Auxin Connection

Long-term (6+ h) auxin responses have been shown in rice to rely on the actin binding protein RMD (Rice Morphology Determinant; FORMIN5, homolog of Arabidopsis FORMIN14), which is downstream of auxin response factors (Zhang et al., 2011; Li et al., 2014a). Less is known about auxin’s effect on actin during cellular activities that occur on the order of minutes such as polarized growth or gravitropism (Xu et al., 2014; Zhu et al., 2015). Substantial membrane depolarization, slight acidification of cytosol (in conjunction with significant alkalinization of extracellular pH), and significant but transient increases in cytosolic Ca^2+^ are short-term intracellular responses that were shown to occur independent of transcriptional responses and are AUX1-dependent (Monshausen et al., 2011; Dindas et al., 2018). Actin binding proteins that are known to be modulated by pH or Ca^2+^, such as ADF/cofilin or villin, would be good candidates for a target of the hormone and mutants could be evaluated for growth in the presence of IAA or a lack of actin reorganization in response to 20–30 min IAA treatments. The drastic auxin-induced actin reorganization could require more than one actin binding protein; perhaps ecotype-specific differences in actin binding protein expression explain Col-0 and WS’s dissimilar responses to NAA at 20–30 min.

### The Actin–Auxin Connection and Cell Expansion

The mechanisms by which auxin and actin control growth are unknown. Actin could function by providing tracks for trafficking auxin transporters, altering vacuole morphology, and/or operating through another mechanism. Auxin efflux carriers (PIN proteins) and the influx protein AUX1 were previously shown to depend on actin for targeted subcellular localization (Kleine-Vehn et al., 2006; 2008). However, AUX1 is not redistributed in response to NAA (Kleine-Vehn, et al., 2006), to which we observe at least some actin reorganization in Col-0, WS, and both *aux1* mutants. Further, use of a photoconvertible PIN2 shows that it is not maintained at the root epidermal cell plasma membrane after auxin treatments, and that most PIN2 in brefeldin A compartments is newly synthesized rather than recycled (Jasik et al., 2016). These results complicate a role for actin in trafficking auxin transporters in response to auxin; however, actin reorganization could provide tracks to transport signaling elements into the nucleus.

Turgor pressure exerted by the vacuole (and a loosened cell wall) is a primary driver of cell expansion (Cosgrove, 2005; Kroeger et al., 2011; Braidwood et al., 2013; Guerriero et al., 2014). Six-hour NAA treatments cause actin-dependent vacuole constriction that ultimately leads to reduced cell lengths (Scheuring et al., 2016). If the same mechanism impels growth cessation within minutes, the denser, more longitudinal actin array we detected might effect vacuole constriction.

Auxin-induced actin reorganization could operate primarily in signaling, and be incidental to changes in growth rate rather than driving growth per se. Whereas wildtype (Col-0) roots ceased elongating within 30 s of low-dose IAA treatments, IAA did not significantly affect *aux1-100* root elongation, and 100 nM NAA reduced root growth rate to approximately wildtype levels (Fendrych et al., 2018). At a similar timepoint after 100 nM NAA treatments, we observed that both *aux1-100* and *aux1-22*, exhibited only partial, and divergent, actin reorganization. If there were direct, causative relationships between actin organization and cell expansion or vice versa, NAA should have induced in *aux1* the complete complement of actin rearrangements observed in wildtype cells or at least the same actin response in both mutant alleles.

Actin reorganization is highly energy intensive, costing as much as 1200 ATP-loaded actin monomers per second during filament elongation (Li et al., 2015b), and even if there is no causal relationship between auxin-induced increased actin density and parallel longitudinality, and cell expansion, it seems unlikely that such extensive reorganization would occur for no functional purpose. It is possible that initial actin reorganization after auxin treatment occurs primarily to transduce the auxin signal. Alternatively, cytoplasmic streaming and vesicle delivery could require an exact equilibrium of available tracks and space in which to move, and any actin array that disrupts that balance quickly alters cell expansion. Toward this idea, Tominaga et al. (2013) showed that faster myosins (i.e., faster delivery along actin tracks) grow larger plants with larger cells, and presumably enhanced cell expansion.

Our IAA treatments provide clear evidence that the actin cytoskeleton in cells along the entire root elongation zone responds to the growth cessation signal within minutes by significantly *increasing* filament abundance as well as, in opposition to the current view that organization leads to or is necessary for expansion, apparent actin organization. We show that IAA-induced actin rearrangements require AUX1, while our NAA results show that auxin itself is able to act as a cytoplasmic signal to modulate actin cytoskeleton organization. We conclude that, however auxin is acting, the relationship between actin organization and cell expansion cannot be explained by a simple model requiring either “organized” or “disorganized” actin, or by a presence or absence of longitudinal bundles.

## METHODS

### Plant Material and Growth Conditions

Roots for all experiments were from 6-day-old, light-grown seedlings expressing GFP-fABD2: Col-0, WS, *aux1-100*, and *aux1-22*. Seeds were surface sterilized and stratified at 4°C for two days. All plants were grown on 0.5× Murashige and Skoog medium solidified with 0.6% (w/v) agar and no sucrose, as described previously (Sheahan et al., 2004; Dyachok et al., 2011; Henty et al., 2011; Li et al., 2014b; Cai et al., 2014). Seedlings were grown at 21°C, vertically and under long-day conditions (16 h of light, 8 h of darkness).

Seeds for *Arabidopsis thaliana* T-DNA insertion mutant *aux1-100* (CS2360) and EMS point mutant *aux1-22* (CS9585) were obtained from the ABRC stock center and, with WS-0 and Col-0, transformed with GFP-fABD2 (Sheahan et al., 2004) using the floral dip method (Zhang et al., 2006). T1 plants were screened on plates with hygromycin. Plants of *aux1-100* were then genotyped by PCR to confirm homozygosity using DNA primers WT-forward 5’-GCATGCTATGTGGAAACCACAGAAG-3’ and WT-reverse 5’-tacCTGACGAGCGGAGGCAGATC-3’ and Feldmann/AZ primers for the mutant: forward 5’-gatgcactcgaaatcagccaattttagac-3’ and reverse 5’-tccttcaatcgttgcggttctgtcagttc-3’. *aux1-22* mutants were identified by their agravitropic phenotype. T2 plants were used for experiments.

### VAEM Imaging, Measuring Cell Lengths, and Quantitative Analysis of Cortical Actin Array Architecture

To measure cell sizes and obtain a corresponding measurement of each actin parameter, we collected overlapping VAEM images (single optical sections) of cortical cytoplasm in root epidermal cells expressing GFP-fABD2. Images were collected from the root apex to the first obviously visible root hair initiations.

VAEM was performed using a TIRF illuminator mounted on an IX-71 microscope equipped with a 60× 1.45–numerical aperture PlanApo TIRF objective (Olympus). Illumination was from a solid-state 50-mW laser (Intelligent Imaging Innovations) attenuated to 3–5% power, depending on the day, but kept the same for a single experiment/replicate. The 488-nm laser emission was captured with an electron multiplying charge-coupled device camera (ORCA-EM C9100-12; Hamamatsu Photonics). The microscope platform was operated and images collected with Slidebook software (version 6; Intelligent Imaging Innovations). A fixed exposure and gain were selected so that individual actin filaments could be seen but higher order filament structures were not intensity-saturated.

Two images were collected per field of view: one to capture actin filaments in focus and one to visualize the cell side and end walls in a higher focal plane, since these are frequently clearly visible in this higher plane without staining. Each image was rotated with an image rotating macro so the longitudinal axes of the cells photographed were parallel to the horizon of the image. All micrographs were cropped and analyzed in FIJI (https://fiji.sc/). For the analysis, we lined up the overlapping images to recreate a full view of the root. In a color (RGB) version of the image stack file, we identified, marked, numbered, and measured cells whose side and end walls were distinguishable, generally choosing cells in the middle of the root to avoid including ones that might present differences in actin architecture due to differences in the cell’s angle relative to the objective. On the RGB image stack, to better distinguish cells, we frequently enhanced brightness and contrast; all cropped images used for quantifying actin architecture and orientation were taken from original 8-bit files. Actin images were cropped along the entire length of every specified cell, and numbered to correspond to the specific cell from which they were cropped. Skewness and density were analyzed according to Higaki et al. (2010b) and Henty et al. (2011); angle and parallelness were analyzed according to Ueda et al. (2010) and Cai et al. (2014). The size of crops must be consistent for all images in an experiment and frequently individual crops were smaller than the entire length of a cell; in such cases an actin measurement was obtained for each crop and the final scatter-plotted measurement for each actin parameter for an individual cell was taken as the mean of the measurements from that cell’s particular set of crops. In Col-0 root characterization, we analyzed cell size and corresponding actin architecture for more than 180 cells from at least 20 roots total—all the cells with clearly distinguishable end walls. For effects of IAA on Col-0, cells up to 85 μm were counted as belonging to “Region 2”, cells more than 94 μm were categorized as “Region 3”, and cells falling between 85–94 μm were counted in both categories. To quantify actin architecture and orientation on a “per cell” basis for the *WS–aux1-100* and *Col-0*–*aux1-22* analyses (Figure 5 and Supplemental Figures 7, 8, and 9), we used the mean value from a single cell’s set of crops as the value representing the actin measurement for that cell. For example, to fully account for all the actin in a 160 μm-long cell, 10 crops would be needed. Measurements on a per cell basis would take the mean of the density values for those 10 crops as a single density value for that cell. In determining *aux1* response to IAA and NAA, we analyzed a minimum of 125 cells (from a total of at least 9 roots) per genotype per treatment. Relationships between actin parameters and cell dimensions were analyzed in Microsoft Excel and JMP.

### Auxin Treatments

IAA was obtained from Sigma-Aldrich (I2886) and diluted to a 10 mM stock concentration in ultrapure ethanol (FisherScientific BP2818500). NAA was also from Sigma-Aldrich (N0640) and diluted to a 10 mM stock concentration in ultrapure ethanol. For experiments, each auxin was further diluted to appropriate concentrations into 0.5× MS liquid medium without sucrose; for mock solution, ultrapure ethanol was added to 0.5× MS liquid medium without sucrose to match the highest concentration of IAA or NAA used. To ensure even IAA or NAA treatment of plants during 20–30 min treatments, whole seedlings were cut from agar plates and treated by soaking on their agar block in a 24-well plate. For the very short-term treatments used for 100-s timelapse movies, plants were treated on slides by being mounted in either mock or IAA solution. Imaging began almost immediately and both regions were imaged within 7 min. For 20–30 min treatments, all imaging concluded within 30 min. Because darkness can stimulate degradation of cytoskeletal organizing proteins (Dyachok et al., 2011) and a reorientation of actin filaments in hypocotyls (Breuer et al., 2014), plants were left under grow lights (while soaking in solution during 20-min treatments) and slides were prepared in the light. All IAA and NAA experiments were performed and analyzed double blind.

### Individual Actin Filament Dynamics

Individual actin filaments were captured with 100-s timelapse VAEM using a 150×1.45 NA UApoN TIRF objective (Olympus). To determine differences in actin filament behavior between shorter and longer cells, we documented cell size by taking snapshots of the entire cells from which the timelapse movies were captured. In general, movies of Region 2 cells were collected from Region 2 cells close to the root cap and movies of Region 3 cells were collected from Region 3 cells close to the end of the elongation zone (i.e., the first cell rootward of the first visible root hair initiation). All timelapse movies and regions of interest were analyzed in FIJI. To best display the representative filaments and their dynamics, brightness and contrast were enhanced in the final montages of Figure 2B and Figure 2C. Occasionally, minimal adjustments to brightness and contrast were made during analysis to more definitively follow some filaments or events. Filament severing frequency, maximum filament length, filament lifetime, and elongation rates were measured as described previously (Staiger et al. 2009; Henty et al., 2011; Cai et al., 2014; Henty-Ridilla et al., 2014). To measure bundling, debundling, and annealing frequencies, we cropped ≈ 15 μm × 15 μm ROIs (exact size 227.7 μm^2^). For measuring individual filament responses to IAA, we used ≈ 7 μm × 7 μm ROIs (exact size 57.8 μm^2^). To account for differences in filament density in short and long cells, bundling, unbundling, and annealing frequencies were normalized against filament numbers in each ROI. An incident of bundling was counted as an incident in which filament fluorescence intensity increased, either from an apparent “catch and zip” event (categorized as a “zippering event”; these events comprise approximately 90% of observed incidents of bundling) or, simply, a visible, unambiguous increase with a minimum three-frame persistence (3 s ≥ 10% filament lifetime) in fluorescence intensity for which “catch and zip” was not specifically apparent (these were categorized as “other bundling event” and account for the remaining ≈ 10% of bundling incidents). Unbundling events were counted as incidents in which a filament was visible next to a mother filament (usually “unpeeling” over several timelapse frames) and, frequently, fluorescence intensity decreased. In cases without a visible decrease in fluorescence intensity, we included only events where the filament clearly “peeled off” from the mother filament. Incidents of annealing were counted when ends of two F-actin fragments joined together for a minimum of two frames. It was not highly unusual to see this annealing behavior join three pieces of recently severed actin filament; if three distinguishable fragments joined to form an individual filament in the same frame, this was counted as two annealing events, one between each fragment.

When capturing timelapse movies to document individual filament changes in response to IAA within 7 min, we applied ≈ 70 μL of either blinded solution (10 nM IAA or mock) directly to the microscope slide, then the root and coverslip, and imaged immediately, alternately imaging Region 2 or Region 3 first so the timepoints of each dataset would average out to 0-7 min from applying the treatment to the slide.

### Accession Numbers

Sequence data from this article can be found in the Arabidopsis Information Resource database (https://www.arabidopsis.org/) under the following names and accession numbers: AUX1 (At2G38120).

## SUPPLEMENTAL MATERIALS

**Supplemental Table 1.** Eigenvectors for Principal Component Analysis of Cell Size vs. Actin Parameters in Col-0.

**Supplemental Table 2.** Eigenvalues for Principal Component Analysis of Cell Size vs. Actin Parameters in Col-0.

**Supplemental Table 3.** Eigenvectors for Principal Component Analysis of Cell Size vs. Actin Parameters in WS.

**Supplemental Table 4.** Eigenvalues for Principal Component Analysis of Cell Size vs. Actin Parameters in WS.

**Supplemental Table 5.** Eigenvectors for Principal Component Analysis of Cell Size vs. Actin Parameters in *aux1*-*100*.

**Supplemental Table 6.** Eigenvalues for Principal Component Analysis of Cell Size vs. Actin Parameters in *aux1*-*100*.

**Supplemental Table 7.** Actin Architecture Measurements after IAA Treatments.

**Supplemental Figure 1.** Epidermal Cells in Different Root Regions Exhibit Distinct Actin Filament Arrays.

**Supplemental Figure 2.** Actin Filament Arrays Are Not Predictive of Cell Width.

**Supplemental Figure 3.** Actin Filament Arrays Are Predictive of Cell Length.

**Supplemental Figure 4.** Actin Arrays in Region 3 Are More Dynamic than in Region 2.

**Supplemental Figure 5.** Short-Term IAA Treatments Induce Dose-Dependent Changes in Actin Filament Organization.

**Supplemental Figure 6.** Short-Term IAA Treatments Induce a Time-Dependent Increase in Actin Filament Density and Longitudinal Orientation.

**Supplemental Figure 7.** Actin Filament Organization Plotted with Respect to Corresponding Cell Length in WS and *aux1-100*.

**Supplemental Figure 8.** Actin Organization in *aux1-22* Fails to Respond to Short-Term IAA Treatments but Partially Responds to the Membrane-Permeable Auxin NAA.

**Supplemental Figure 9.** Actin Filament Organization Plotted with Respect to Corresponding Cell Length in Col-0 and *aux1-22*.

**Supplemental Figure 10.** Hypothetical model of auxin perception by AUX1 upstream of actin cytoskeleton reorganization.

## ACKNOWLEDGEMENTS

This work was supported, in part, by an award from the Office of Science at the US Department of Energy, Physical Biosciences Program, under contract number DE-FG02-09ER15526 to C.J.S.

## AUTHOR CONTRIBUTIONS

R.S.A and C.J.S conceived the project and designed the experiments; R.S.A performed the experiments and data analysis; and R.S.A and C.J.S. wrote the article.

## REFERENCES

Andrianantoandro, E., Blanchoin, L., Sept, D., McCammon, J.A., and Pollard, T.D. (2001). Kinetic mechanism of end-to-end annealing of actin filaments. J. Mol. Biol. 312: 721–730. doi:10.1006/jmbi.2001.5005.

Baluška, F., Jasik, J., Edelmann, H.G., Salajová, T., and Volkmann, D. (2001). Latrunculin B-induced plant dwarfism: plant cell elongation is F-actin-dependent. Dev. Biol. 231: 113–124. doi:10.1006/dbio.2000.0115.

Baluška, F., and Mancuso, S. (2013). Root apex transition zone as oscillatory zone. Front. Plant Sci. 4: e354. doi:10.3389/fpls.2013.00354.

Baluška, F., Vitha, S., Barlow, P.W., and Volkmann, D. (1997). Rearrangements of F-actin arrays in growing cells of intact maize root apex tissues: a major developmental switch occurs in the postmitotic transition region. Eur. J. Cell Biol. 72: 113–121.

Band, L.R., Wells, D.M., Larrieu, A., Sun, J., Middleton, A.M., French, A.P., Brunoud, G., Sato, E.M., Wilson, M.H., Péret, B., Oliva, M., Swarup, R., Sairanen, I., Parry, G., Ljung, K., Beeckman, T., Garibaldi, J.M., Estelle, M., Owen, M.R., Vissenberg, K., Hodgman, T.C., Pridmore, T.P., King, J.R., Vernoux, T., and Bennett, M.J. (2012). Root gravitropism is regulated by a transient lateral auxin gradient controlled by a tipping-point mechanism. Proc. Natl. Acad. Sci. USA 109: 4668–4673. doi:10.1073/pnas.1201498109.

Beemster, G.T.S., and Baskin, T.I. (1998). Analysis of cell division and elongation underlying the developmental acceleration of root growth in *Arabidopsis thaliana*. Plant Physiol. 116: 1515–1526. doi:10.1104/pp.116.4.1515.

Beemster, G.T.S., and Baskin, T.I. (2000). *STUNTED PLANT 1* mediates effects of cytokinin, but not of auxin, on cell division and expansion in the root of Arabidopsis. Plant Physiol. 124: 1718–1727. doi:10.1104/pp.124.4.1718.

Bennett, M.J., Marchant, A., Green, H.G., May, S.T., Ward, S.P., Millner, P.A., Walker, A.R., Schulz, B., and Feldmann, K.A. (1996). *Arabidopsis AUX1* gene: a permease-like regulator of root gravitropism. Science 273: 948–950. doi:10.1126/science.273.5277.948.

Braidwood, L., Breuer, C., and Sugimoto, K. (2014). My body is a cage: Mechanisms and modulation of plant cell growth. New Phytol. 201: 388–402. doi:10.1111/nph.12473.

Breuer, D., Ivakov, A., Sampathkumar, A., Hollandt, F., Persson, S., and Nikoloski, Z. (2014). Quantitative analyses of the plant cytoskeleton reveal underlying organizational principles. J. Royal Soc. Interface 11: e20140362. doi:10.1098/rsif.2014.0362.

Brunoud, G., Wells, D. M., Oliva, M., Larrieu, A., Mirabet, V., Burrow, A. H., Beeckman, T., Kepinski, S., Traas, J., Bennett, M.J., Vernoux, T. (2012). A novel sensor to map auxin response and distribution at high spatio-temporal resolution. Nature 482: 103–106. doi:10.1038/nature10791.

Cai, C., Henty-Ridilla, J.L., Szymanski, D.B., and Staiger, C.J. (2014). Arabidopsis myosin XI: A motor rules the tracks. Plant Physiol. 166: 1359–1370. doi:10.1104/pp.114.244335.

Calderón Villalobos, L.I.A., Lee, S., De Oliveira, C., Ivetac, A., Brandt, W., Armitage, L., Sheard, L. B., Tan, X., Parry, G., Mao, H., Zheng, N., Napier, R., Kepinski, S., and Estelle, M. (2012). A combinatorial TIR1/AFB-Aux/IAA coreceptor system for differential sensing of auxin. Nat. Chem. Biol 8: 477–485. doi:10.1038/nchembio.926.

Cao, L., Henty-Ridilla, J.L., Blanchoin, L., and Staiger, C.J. (2016). Profilin-dependent nucleation and assembly of actin filaments controls cell elongation in Arabidopsis. Plant Physiol. 170: 220–233. doi:10.1104/pp.15.01321.

Cárdenas, L., Vidali, L., Domínguez, J., Pérez, H., Sánchez, F., Hepler, P.K., and Quinto, C. (1998). Rearrangement of actin microfilaments in plant root hairs responding to rhizobium ETLI nodulation signals. Plant Physiol. 116: 871–877. doi:10.1104/pp.116.3.871.

Carrier, D.J., Bakar, N.T.A., Swarup, R., Callaghan, R., Napier, R.M., Bennett, M.J., and Kerr, I.D. (2008). The binding of auxin to the Arabidopsis auxin influx transporter AUX1. Plant Physiol. 148: 529–535. doi:10.1104/pp.108.122044.

Chen, J.-G, Ullah, H., Young, J. C., Sussman, M.R., and Jones, A.M. (2001). ABP1 is required for organized cell elongation and division in *Arabidopsis* embryogenesis. Genes Dev. 15: 902–911. doi:10.1101/gad.866201.

Chen, X., Naramoto, S., Robert, S., Tejos, R., Löfke, C., Lin, D., Yang, Z., and Friml, J. (2012). ABP1 and ROP6 GTPase signaling regulate clathrin-mediated endocytosis in *Arabidopsis* roots. Curr. Biol. 22: 1326–1332. doi:10.1016/j.cub.2012.05.020.

Cosgrove, D.J. (2005). Growth of the plant cell wall. Nat. Rev. Mol. Cell Biol. 6: 850–861. doi:10.1038/nrm1746.

Dai, X., Zhang, Y., Zhang, D., Chen, J., Gao, X., Estelle, M., and Zhao, Y. (2015). Embryonic lethality of *Arabidopsis abp1-1* is caused by deletion of the adjacent BSM gene. Nat. Plants 1: e15183. doi:10.1038/nplants.2015.183.

Delker, C., Pöschl, Y., Raschke, A., Ullrich, K., Ettingshausen, S., Hauptmann, V., Grosse, I., and Quint, M. (2010). Natural variation of transcriptional auxin response networks in *Arabidopsis thaliana*. Plant Cell 22: 2184–2200. doi:10.1105/tpc.110.073957.

Dharmasiri, N., Dharmasiri, S., and Estelle, M. (2005a). The F-box protein TIR1 is an auxin receptor. Nature 435: 441–445. doi:10.1038/nature03543.

Dharmasiri, N., Dharmasiri, S., Weijers, D., Lechner, E., Yamada, M., Hobbie, L., Ehrismann, J.S., Jürgens, G., and Estelle, M. (2005b). Plant development is regulated by a family of auxin receptor F-box proteins. Dev. Cell 9: 109–119. doi:10.1016/j.devcel.2005.05.014.

Dhonukshe, P., Grigoriev, I., Fischer, R., Tominaga, M., Robinson, D. G., Hašek, J., Paciorek, T., Petrášek, J., Seifertová, D., Tejos, R., Meisel, L.A., Zažímalová, E., Gadella, Jr., T.W.J., Steirhof, Y.-D., Ueda, T., Oiwa, K., Akhmanova, A., Brock, R., Spang, A., and Friml, J. (2008). Auxin transport inhibitors impair vesicle motility and actin cytoskeleton dynamics in diverse eukaryotes. Proc. Natl. Acad. Sci. USA 105: 4489–4494. doi:10.1073/pnas.0711414105.

Dindas, J., Scherzer, S., Roelfsema, M.R.G., Von Meyer, K., Müller, H.M., Al-Rasheid, K.A.S., Palme, K., Dietrich, P., Becker, D., Bennett, M.J., and Hedrich, R. (2018). AUX1-mediated root hair auxin influx governs SCFTIR1/AFB-type Ca^2+^ signaling. Nat. Commun. 9: e1174. doi:10.1038/s41467-018-03582-5.

Dyachok, J., Zhu, L., Liao, F., He, J., Huq, E., and Blancaflor, E.B. (2011). SCAR mediates light-induced root elongation in *Arabidopsis* through photoreceptors and proteasomes. Plant Cell 23: 3610–3626. doi:10.1105/tpc.111.088823.

Falasca, G., and Altamura, M.M. (2003). Histological analysis of adventitious rooting in *Arabidopsis thaliana* (L.) Heynh seedlings. Plant Biosyst. 137: 265–273. dx.doi.org/10.1080/11263500312331351511.

Feldmann, K.A. (1991). T-DNA insertion mutagenesis in *Arabidopsis*: Mutational spectrum. Plant J. 1: 71–82. doi:10.1111/j.1365-313X.1991.00071.x.

Fendrych, M., Akhmanova, M., Merrin, J., Glanc, M., Hagihara, S., Takahashi, K., Uchida, N., Torii, K.U., and Friml, J. (2018). Rapid and reversible root growth inhibition by TIR1 auxin signalling. Nat. Plants 4: 453–459. doi:10.1038/s41477-018-0190-1.

Gao, Y., Zhang, Y., Zhang, D., Dai, X., Estelle, M., and Zhao, Y. (2015). AUXIN BINDING PROTEIN 1 (ABP1) is not required for either auxin signaling or *Arabidopsis* development. Proc. Natl. Acad. Sci. USA 112: 2275–2280. doi:10.1073/pnas.1500365112.

Gilliland, L.U., Pawloski, L.C., Kandasamy, M.K., and Meagher, R.B. (2003). *Arabidopsis* actin gene *ACT7* plays an essential role in germination and root growth. Plant J. 33: 319–328. doi:10.1046/j.1365-313X.2003.01626.x.

Grones, P. and Friml, J. (2015). Auxin transporters and binding proteins at a glance. J. Cell Sci. 128: 1–7. doi:10.1242/jcs.159418.

Guerriero, G., Hausman, J. -F, and Cai, G. (2014). No stress! Relax! Mechanisms governing growth and shape in plant cells. Int. J. Mol. Sci. 15: 5094–5114. doi:10.3390/ijms15035094.

Hacham, Y., Holland, N., Butterfield, C., Ubeda-Tomas, S., Bennett, M.J., Chory, J., and Savaldi-Goldstein, S. (2011). Brassinosteroid perception in the epidermis controls root meristem size. Development 138: 839–848. doi:10.1242/dev.061804.

Hayashi, K.-I, Nakamura, S., Fukunaga, S., Nishimura, T., Jenness, M.K., Murphy, A.S., Motose, H., Nozaki, H., Furutani, M., and Aoyama, T. (2014). Auxin transport sites are visualized in planta using fluorescent auxin analogs. Proc. Natl. Acad. Sci. USA 111: 11557–11562. doi:10.1073/pnas.1408960111.

Hejnowicz, Z., and Erickson, R.O. (1968). Growth inhibition and recovery in roots following temporary treatment with auxin. Physiol. Plantarum 21: 302–313. doi:10.1111/j.1399-3054.1968.tb07254.x.

Henty, J.L., Bledsoe, S.W., Khurana, P., Meagher, R.B., Day, B., Blanchoin, L., and Staiger, C.J. (2011). *Arabidopsis* ACTIN DEPOLYMERIZING FACTOR4 modulates the stochastic dynamic behavior of actin filaments in the cortical array of epidermal cells. Plant Cell 23: 3711–3726. doi:10.1105/tpc.111.090670.

Henty-Ridilla, J.L., Li, J., Blanchoin, L., and Staiger, C.J. (2013). Actin dynamics in the cortical array of plant cells. Curr. Opin. Plant Biol. 16: 678–687. doi:10.1016/j.pbi.2013.10.012.

Henty-Ridilla, J.L., Li, J., Day, B., and Staiger, C.J. (2014). ACTIN DEPOLYMERIZING FACTOR4 Regulates actin dynamics during innate immune signaling in *Arabidopsis*. Plant Cell 26: 340–352. doi:10.1105/tpc.113.122499.

Higaki, T., Kojo, K.H., and Hasezawa, S. (2010a.) Critical role of actin bundling in plant cell morphogenesis. Plant Signal. Behav. 5: 484–488. doi:10.4161/psb.10947.

Higaki, T., Kurusu, T., Hasezawa, S., and Kuchitsu, K. (2011). Dynamic intracellular reorganization of cytoskeletons and the vacuole in defense responses and hypersensitive cell death in plants. J. Plant Res. 124: 315–324. doi:10.1007/s10265-011-0408-z.

Higaki, T., Kutsuna, N., Sano, T., Kondo, N., and Hasezawa, S. (2010b.) Quantification and cluster analysis of actin cytoskeletal structures in plant cells: role of actin bundling in stomatal movement during diurnal cycles in Arabidopsis guard cells. Plant J. 61: 156–165. doi:10.1111/j.1365-313X.2009.04032.x.

Holweg, C., Süßlin, C., and Nick, P. (2004). Capturing in vivo dynamics of the actin cytoskeleton stimulated by auxin or light. Plant Cell. Physiol. 45: 855–863. doi:10.1093/pcp/pch102.

Hussey, P.J., Ketelaar, T., and Deeks, M.J. (2006). Control of the actin cytoskeleton in plant cell growth. Annu. Rev. Plant Biol. 57: 109–125. doi:10.1146/annurev.arplant.57.032905.105206.

Jásik, J., Bokor, B., Stuchlík, S., Mičieta, K., Turňa, J., and Schmelzer, E. (2016). Effects of auxins on PIN-FORMED2 (PIN2) dynamics are not mediated by inhibiting PIN2 endocytosis. Plant Physiol. 172: 1019–1031. doi:10.1104/pp.16.00563.

Kandasamy, M.K., McKinney, E.C., and Meagher, R.B. (2009). A single vegetative actin isovariant overexpressed under the control of multiple regulatory sequences is sufficient for normal *Arabidopsis* development. Plant Cell 21: 701–718. doi:10.1105/tpc.108.061960.

Kepinski, S. and Leyser, O. (2005). The *Arabidopsis* F-box protein TIR1 is an auxin receptor. Nature 435: 446–451. doi:10.1038/nature03542.

Ketelaar, T., De Ruijter, N.C.A., and Emons, A.M.C. (2003). Unstable F-actin specifies the area and microtubule direction of cell expansion in Arabidopsis root hairs. Plant Cell 15: 285–292. doi:10.1105/tpc.007039.

Kleine-Vehn, J., Dhonukshe, P., Sauer, M., Brewer, P.B., Wiśniewska, J., Paciorek, T., Benková, E., and Friml, J. (2008). ARF GEF-dependent transcytosis and polar delivery of PIN auxin carriers in *Arabidopsis*. Curr. Biol. 18: 526–531. doi:10.1016/j.cub.2008.03.021.

Kleine-Vehn, J., Dhonukshe, P., Swarup, R., Bennett, M., and Friml, J. (2006). Subcellular trafficking of the *Arabidopsis* auxin influx carrier AUX1 uses a novel pathway distinct from PIN1. Plant Cell 18: 3171–3181. doi:10.1105/tpc.106.042770.

Kleine-Vehn, J. and Friml, J. (2008). Polar targeting and endocytic recycling in auxin-dependent plant development. Annu. Rev. Cell Dev. Biol. 24: 447–473. doi:10.1146/annurev.cellbio.24.110707.175254.

Kroeger, J.H., Zerzour, R., and Geitmann, A. (2011). Regulator or driving force? The role of turgor pressure in oscillatory plant cell growth. PLoS ONE 6: e18549. doi:10.1371/journal.pone.0018549.

Leucci, M.R., Di Sansebastiano, G. -P, Gigante, M., Dalessandro, G., and Piro, G. (2007). Secretion marker proteins and cell-wall polysaccharides move through different secretory pathways. Planta 225: 1001–1017. doi:10.1007/s00425-006-0407-9.

Li, G., Liang, W., Zhang, X., Ren, H., Hu, J., Bennett, M.J., and Zhang, D. (2014a). Rice actin-binding protein RMD is a key link in the auxin-actin regulatory loop that controls cell growth. Proc. Natl. Acad. Sci. USA 111: 10377–10382. doi:10.1073/pnas.1401680111.

Li, J., Arieti, R., and Staiger, C.J. (2015a). “Actin filament dynamics and their role in plant cell expansion.” pp. 127–162 in Plant Cell Wall Patterning and Cell Shape, H. Fukuda, ed. doi:10.1002/9781118647363.ch5.

Li, J., Henty-Ridilla, J.L., Huang, S., Wang, X., Blanchoin, L., and Staiger, C.J. (2012). Capping protein modulates the dynamic behavior of actin filaments in response to phosphatidic acid in *Arabidopsis*. Plant Cell 24: 3742–3754. doi:10.1105/tpc.112.103945.

Li, J., Henty-Ridilla, J.L., Staiger, B.H., Day, B., and Staiger, C.J. (2015b). Capping protein integrates multiple MAMP signalling pathways to modulate actin dynamics during plant innate immunity. Nat. Commun. 6: 7206. doi:10.1038/ncomms8206.

Li, J., Staiger, B.H., Henty-Ridilla, J.L., Abu-Abied, M., Sadot, E., Blanchoin, L., and Staiger, C.J. (2014b). The availability of filament ends modulates actin stochastic dynamics in live plant cells. Mol. Biol. Cell 25: 1263–1275. doi:10.1091/mbc.E13-07-0378.

Lin, D., Nagawa, S., Chen, J., Cao, L., Chen, X., Xu, T., Li, H., Dhonukshe, P., Yamamuro, C., Friml, J., Scheres, B., Fu, Y., Yang, Z. (2012). A ROP GTPase-dependent auxin signaling pathway regulates the subcellular distribution of PIN2 in *Arabidopsis* roots. Curr. Biol. 22: 1319–1325. doi:10.1016/j.cub.2012.05.019.

Maher, E.P., and Martindale, S.J.B. (1980). Mutants of *Arabidopsis thaliana* with altered responses to auxins and gravity. Biochem. Genet. 18: 1041–1053. doi:10.1007/BF00484337.

Marchant, A., Kargul, J., May, S.T., Muller, P., Delbarre, A., Perrot-Rechenmann, C., and Bennett, M.J. (1999). AUX1 regulates root gravitropism in *Arabidopsis* by facilitating auxin uptake within root apical tissues. EMBO J. 18: 2066–2073. doi:10.1093/emboj/18.8.2066.

Mathur, J. (2004). Cell shape development in plants. Trends Plant Sci. 9: 583–590. doi:10.1016/j.tplants.2004.10.006.

Monshausen, G.B., Miller, N.D., Murphy, A.S., and Gilroy, S. (2011). Dynamics of auxin-dependent Ca^2+^ and pH signaling in root growth revealed by integrating high-resolution imaging with automated computer vision-based analysis. Plant J. 65: 309–318. doi:10.1111/j.1365-313X.2010.04423.x.

Nagawa, S., Xu, T., Lin, D., Dhonukshe, P., Zhang, X., Friml, J., Scheres, B., Fu, Y., and Yang, Z. (2012). ROP GTPase-dependent actin microfilaments promote PIN1 polarization by localized inhibition of clathrin-dependent endocytosis. PLoS Biol. 10: e1001299. doi:10.1371/journal.pbio.1001299.

Nick, P. (2010). Probing the actin-auxin oscillator. Plant Signal. Behav. 5: 94–98. doi:10.4161/psb.5.2.10337.

Nick, P., Han, M.-J, and An, G. (2009). Auxin stimulates its own transport by shaping actin filaments. Plant Physiol. 151: 155–167. doi:10.1104/pp.109.140111.

Overvoorde, P., Fukaki, H., and Beeckman, T. (2010). Auxin control of root development. CSH Perspect. Biol. 2: a001537. doi:10.1101/cshperspect.a001537.

Pickett, F.B., Wilson, A.K., and Estelle, M. (1990). The *aux1* mutation of *Arabidopsis* confers both auxin and ethylene resistance. Plant Physiol. 94: 1462–1466. doi:10.1104/pp.94.3.1462.

Rahman, A., Bannigan, A., Sulaman, W., Pechter, P., Blancaflor, E.B., and Baskin, T.I. (2007). Auxin, actin and growth of the *Arabidopsis thaliana* primary root. Plant J. 50: 514–528. doi:10.1111/j.1365-313X.2007.03068.x.

Rashotte, A.M., Poupart, J., Waddell, C.S., and Muday, G.K. (2003). Transport of the two natural auxins, indole-3-butyric acid and indole-3-acetic acid, in Arabidopsis. Plant Physiol. 133: 761–772. doi:10.1104/pp.103.022582.

Roman G, Lubarsky, B., Kieber, J.J., Rothenberg, M., and Ecker, J.R. (1995). Genetic analysis of ethylene signal transduction in *Arabidopsis thaliana*: Five novel mutant loci integrated into a stress response pathway. Genetics 139: 1393–1409.

Rutschow, H.L., Baskin, T.I., and Kramer, E.M. (2014). The carrier AUXIN RESISTANT (AUX1) dominates auxin flux into Arabidopsis protoplasts. New Phytol. 204: 536–544. doi:10.1111/nph.12933.

Sauer, M., and Kleine-Vehn, J. (2011). AUXIN BINDING PROTEIN1: The outsider. Plant Cell 23: 2033–2043. doi:10.1105/tpc.111.087064.

Savaldi-Goldstein, S., Peto, C., and Chory, J. (2007). The epidermis both drives and restricts plant shoot growth. Nature 446: 199–202. doi:10.1038/nature05618.

Scheuring, D., Löfke, C., Krüger, F., Kittelmann, M., Eisa, A., Hughes, L., Smith, R.S., Hawes, C., Schumacher, K., and Kleine-Vehn, J. (2016). Actin-dependent vacuolar occupancy of the cell determines auxin-induced growth repression. Proc. Natl. Acad. Sci. USA 113: 452–457. doi:10.1073/pnas.1517445113.

Schultz, E.R., Zupanska, A.K., Sng, N.J., Paoul, A-L., and Ferl, R.J. (2017). Skewing in Arabidopsis roots involves disparate environmental signaling pathways. BMC Plant Biol. 17: 31. doi:10.1186/s12870-017-0975-9.

Sheahan, M.B., Staiger, C.J., Rose, R.J., and McCurdy, D.W. (2004). A green fluorescent protein fusion to actin-binding domain 2 of Arabidopsis fimbrin highlights new features of a dynamic actin cytoskeleton in live plant cells. Plant Physiol. 136: 3968–3978. doi:10.1104/pp.104.049411.

Smertenko, A.P., Deeks, M.J., and Hussey, P.J. (2010). Strategies of actin reorganisation in plant cells. J. Cell Sci. 123: 3019–3028. doi:10.1242/jcs.071126.

Staiger, C.J., Sheahan, M.B., Khurana, P., Wang, X., McCurdy, D.W., and Blanchoin, L. (2009). Actin filament dynamics are dominated by rapid growth and severing activity in the Arabidopsis cortical array. J. Cell Biol. 184: 269–280. doi:10.1083/jcb.200806185.

Swarup, R., Kargul, J., Marchant, A., Zadik, D., Rahman, A., Mills, R., Yemm, A., May, S., Williams, L., Millner, P., Tsurumi, S., Moore, I., Napier, R., Kerr, I.D., Bennett, M.J. (2004). Structure-function analysis of the presumptive Arabidopsis auxin permease AUX1. Plant Cell 16: 3069–3083. doi:10.1105/tpc.104.024737.

Swarup, R., and Péret, B. (2012). AUX/LAX Family of auxin influx carriers-an overview. Front. Plant Sci. 3: e225. doi:10.3389/fpls.2012.00225.

Szymanski, D.B., and Cosgrove, D.J. (2009). Dynamic coordination of cytoskeletal and cell wall systems during plant cell morphogenesis. Curr. Biol. 19: R800–R811. doi:10.1016/j.cub.2009.07.056.

Szymanski, D., and Staiger, C.J. (2017). The actin cytoskeleton: functional arrays for cytoplasmic organization and cell shape control. Plant Physiol. 176: 106–118. doi:https://doi.org/10.1104/pp.17.01519.

Thomas, C. (2012). Bundling Actin filaments from membranes: Some novel players. Front. Plant Sci. 3: e188. doi:10.3389/fpls.2012.00188.

Tominaga, M., Kimura, A., Yokota, E., Haraguchi, T., Shimmen, T., Yamamoto, K., Nakano, A., and Ito, K. (2013). Cytoplasmic streaming velocity as a plant size determinant. Dev. Cell 27: 345–352. doi:10.1016/j.devcel.2013.10.005.

Ueda, H., Yokota, E., Kutsuna, N., Shimada, T., Tamura, K., Shimmen, T., Hasezawa, S., Dolja, V. V., and Hara-Nishimuraa, I. (2010). Myosin-dependent endoplasmic reticulum motility and F-actin organization in plant cells. Proc. Natl. Acad. Sci. USA 107: 6894–6899. doi:10.1073/pnas.0911482107.

Ugartechea-Chirino, Y., Swarup, R., Swarup, K., Péret, B., Whitworth, M., Bennett, M., and Bougourd, S. (2010). The AUX1 LAX family of auxin influx carriers is required for the establishment of embryonic root cell organization in *Arabidopsis thaliana*. Ann. Bot. 105: 277–289. doi:10.1093/aob/mcp287.

Ulmasov, T., Hagen, G., and Guilfoyle, T.J. (1999). Activation and repression of transcription by auxin-response factors. Proc. Natl. Acad. Sci. USA 96: 5844–5849. doi:10.1073/pnas.96.10.5844.

van der Weele, C.M., Jiang, H.S., Palaniappan, K.K., Ivanov, V.B., Palaniappan, K., and Baskin, T.I. (2003). A new algorithm for computational image analysis of deformable motion at high spatial and temporal resolution applied to root growth. roughly uniform elongation in the meristem and also, after an abrupt acceleration, in the elongation zone. Plant Physiol. 132: 1138–1148. doi:10.1104/pp.103.021345.

Vidali, L., Burkart, G.M., Augustine, R.C., Kerdavid, E., Tüzel, E., and Bezanilla, M. (2010). Myosin XI is essential for tip growth in *Physcomitrella patens*. Plant Cell 22: 1868–1882. doi:10.1105/tpc.109.073288.

Wu, S., Xie, Y., Zhang, J., Ren, Y., Zhang, X., Wang, J., Guo, X.,Wu, F., Sheng, P., Wang, J., Wu, C., Wang, H., Huang, S., and Wan, J. (2015). VLN2 regulates plant architecture by affecting microfilament dynamics and polar auxin transport in rice. Plant Cell 27: 2829–2845. doi:10.1105/tpc.15.00581.

Xu, T., Dai, N., Chen, J., Nagawa, S., Cao, M., Li, H., Zhou, Z., Chen, X., De Rycke, R., Rakusová, H., Wang, W., Jones, A.M., Friml, J., Patterson, S.E., Bleecker, A.B., Yang, Z. (2014). Cell surface ABP1-TMK auxin-sensing complex activates ROP GTPase signaling. Science 343: 1025–1028. doi:10.1126/science.1245125.

Xu, T., Nagawa, S., and Yang, Z. (2011). Uniform auxin triggers the Rho GTPase-dependent formation of interdigitation patterns in pavement cells. Small GTPases 2: 227–232. doi:10.4161/sgtp.2.4.16702.

Xu, T., Wen, M., Nagawa, S., Fu, Y., Chen, J. -G, Wu, M. -J, Perrot-Rechenmann, C., Friml, J., Jones, A.M., and Yang, Z. (2010). Cell surface- and Rho GTPase-based auxin signaling controls cellular interdigitation in *Arabidopsis*. Cell 143: 99–110. doi:10.1016/j.cell.2010.09.003.

Yanagisawa, M., Desyatova, A.S., Belteton, S.A., Mallery, E.L., Turner, J.A., and Szymanski, D.B. (2015). Patterning mechanisms of cytoskeletal and cell wall systems during leaf trichome morphogenesis. Nat. Plants 1: e 15014. doi:10.1038/nplants.2015.14.

Yang, W., Ren, S., Zhang, X., Gao, M., Ye, S., Qi, Y., Zheng, Y., Wang, J., Zeng, L., Li, Q., Huang, S., and He, Z. (2011). BENT UPPERMOST INTERNODE1 encodes the class II formin FH5 crucial for actin organization and rice development. Plant Cell 23: 661–680. doi:10.1105/tpc.110.081802.

Yang, Y., Hammes, U.Z., Taylor, C. G., Schachtman, D.P., and Nielsen, E. (2006). High-affinity auxin transport by the AUX1 influx carrier protein. Curr. Biol. 16: 1160. doi:10.1016/j.cub.2006.05.043.

Yoshimitsu, Y., Tanaka, K., Fukuda, W., Asami, T., Yoshida, S., Hayashi, K.-i, Kamiya, Y., Jikumaru, Y., Shigeta, T., Nakamura, Y., Matsuo, T., and Okamoto, S. (2011). Transcription of *DWARF4* plays a crucial role in auxin-regulated root elongation in addition to brassinosteroid homeostasis in *Arabidopsis thaliana*. PLoS ONE 6: e23851. doi:10.1371/journal.pone.0023851.

Zhang, W., Cai, C., and Staiger, C.J. (2019). Myosins XI are involved in exocytosis of cellulose synthase complexes. Plant Physiol. 179: 1537–1555. doi:10.1104/pp.19.00018.

Zhang, X., Henriques, R., Lin, S-S., Niu, Q.W., and Chua, N-H. (2006). Agrobacterium-mediated transformation of *Arabidopsis thaliana* using the floral dip method. Nature Protoc. 1: 641–646. doi:10.1038/nprot.2006.97.

Zhang, Z., Zhang, Y., Tan, H., Wang, Y., Li, G., Liang, W., Yuan, Z., Hu, J., Ren, H., and Zhang, D. (2011). RICE MORPHOLOGY DETERMINANT encodes the type II formin FH5 and regulates rice morphogenesis. Plant Cell 23: 681–700. doi:10.1105/tpc.110.081349.

Zhu, J., Bailly, A., Zwiewka, M., Sovero, V., Di Donato, M., Ge, P., Oehri, J., Aryal, B., Hao, P., Linnert, M., Burgardt, N.I., Lücke, C., Weiwad, M., Michel, M., Weiergräber, O.H., Pollmann, S., Azzarello, E., Mancuso, S., Ferro, N., Fukao, Y., Hoffmann, C., Wedlich-Söldner, R., Friml, J., Thomas, C., and Geisler, M. (2016). TWISTED DWARF1 mediates the action of auxin transport inhibitors on actin cytoskeleton dynamics. Plant Cell 28: 930–948. doi:10.1105/tpc.15.00726.

Zhu, J., and Geisler, M. (2015). Keeping it all together: Auxin-actin crosstalk in plant development. J. Exp. Bot. 66: 4983–4998. doi:10.1093/jxb/erv308.

