## Supplemental Tables and Figures for "Auxin-Induced Actin Cytoskeleton Rearrangements Require AUX1"

**Supplemental Table 1.** Eigenvectors for Principal Component Analysis of Cell Size vs. Actin Parameters in Col-0

| Item | Prin1 | Prin2 | Prin3 | Prin4 | Prin5 | Prin6 |
|---|---|---|---|---|---|---|
| Cell Length | 0.527 | 0.093 | 0.048 | 0.044 | 0.609 | -0.582 |
| Cell Width | -0.032 | 0.876 | -0.392 | -0.270 | -0.069 | -0.014 |
| Density | -0.360 | 0.096 | -0.361 | 0.813 | 0.266 | -0.000 |
| Skewness | 0.440 | 0.238 | 0.273 | 0.495 | -0.634 | -0.168 |
| Angle | -0.325 | 0.392 | 0.797 | 0.072 | 0.284 | 0.137 |
| Parallelness | 0.542 | 0.067 | -0.053 | 0.122 | 0.266 | 0.784 |

**Supplemental Table 2.** Eigenvalues for Principal Component Analysis of Cell Size vs. Actin Parameters in Col-0

| Number | Eigenvalue | Percent | Cum. Percent |
|---|---|---|---|
| 1 | 2.9322 | 48.871 | 48.871 |
| 2 | 1.0720 | 17.867 | 66.737 |
| 3 | 0.7749 | 12.914 | 79.652 |
| 4 | 0.7373 | 12.288 | 91.940 |
| 5 | 0.3217 | 5.362 | 97.302 |
| 6 | 0.1619 | 2.698 | 100.000 |

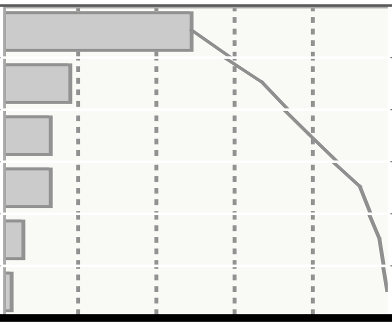

**Supplemental Table 3.** Eigenvectors for Principal Component Analysis of Cell Size vs. Actin Parameters in WS

| Item | Prin1 | Prin2 | Prin3 | Prin4 | Prin5 | Prin6 |
|---|---|---|---|---|---|---|
| WS Cell Length | 0.53137 | 0.13361 | 0.01256 | 0.10846 | 0.34014 | 0.75643 |
| WS Cell Width | -0.02484 | 0.91905 | 0.18593 | -0.33084 | 0.00615 | -0.10329 |
| WS Density | -0.33979 | 0.06901 | 0.72268 | 0.57098 | -0.06879 | 0.16357 |
| WS Skewness | 0.48014 | 0.14062 | -0.09248 | 0.30058 | -0.80565 | -0.04142 |
| WS Angle | -0.32848 | 0.33435 | -0.64176 | 0.58653 | 0.15359 | 0.02918 |
| WS Parallelness | 0.51297 | 0.03429 | 0.15030 | 0.34408 | 0.45483 | -0.62276 |

**Supplemental Table 4.** Eigenvalues for Principal Component Analysis of Cell Size vs. Actin Parameters in WS

| Number | Eigenvalue | Percent | Cum. Percent |
|---|---|---|---|
| 1 | 2.9778 | 49.630 | 49.630 |
| 2 | 1.0810 | 18.016 | 67.646 |
| 3 | 0.9457 | 15.762 | 83.408 |
| 4 | 0.4634 | 7.724 | 91.132 |
| 5 | 0.3727 | 6.212 | 97.344 |
| 6 | 0.1594 | 2.656 | 100.000 |

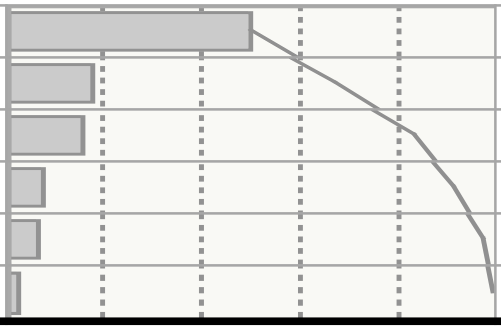

**Supplemental Table 5.** Eigenvectors for Principal Component Analysis of Cell Size vs. Actin Parameters in *aux1-100*

| Item | Prin1 | Prin2 | Prin3 | Prin4 | Prin5 | Prin6 |
|---|---|---|---|---|---|---|
| *aux1-100* Cell Length | 0.47215 | -0.16402 | 0.13255 | 0.22737 | 0.62559 | -0.53810 |
| *aux1-100* Cell Width | 0.21551 | 0.94029 | -0.12556 | -0.17605 | 0.14714 | -0.03177 |
| *aux1-100* Density | -0.39119 | 0.11131 | -0.60851 | 0.66161 | 0.10752 | -0.12251 |
| *aux1-100* Skewness | 0.45730 | 0.10303 | 0.14257 | 0.45766 | -0.70733 | -0.22399 |
| *aux1-100* Angle | -0.35628 | 0.24170 | 0.74402 | 0.43920 | 0.17873 | 0.19035 |
| *aux1-100* Parallelness | 0.49148 | -0.08679 | -0.14993 | 0.27792 | 0.20778 | 0.77976 |

**Supplemental Table 6.** Eigenvalues for Principal Component Analysis of Cell Size vs. Actin Parameters in *aux1-100*

| Number | Eigenvalue | Percent | Cum. Percent |
|---|---|---|---|
| 1 | 3.4137 | 56.895 | 56.895 |
| 2 | 0.9155 | 15.258 | 72.153 |
| 3 | 0.7524 | 12.539 | 84.692 |
| 4 | 0.4141 | 6.901 | 91.593 |
| 5 | 0.3311 | 5.519 | 97.112 |
| 6 | 0.1733 | 2.888 | 100.000 |

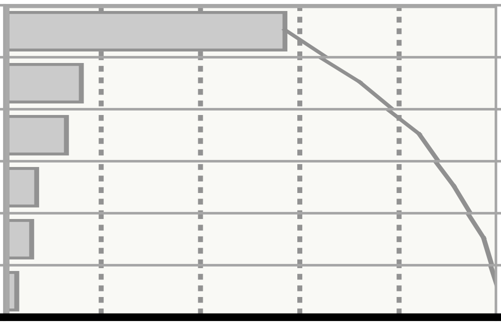

**Supplemental Table 7.** Actin Architecture Measurements after IAA Treatments

| | Region 2 | | | | Region 3 | | | |
|---|---|---|---|---|---|---|---|---|
| **Time** | **Density (%)** | **Skewness** | **Angle (°)** | **Parallelness** | **Density (%)** | **Skewness** | **Angle (°)** | **Parallelness** |
| **≤ 5 min**[1] | 67.9 ± 2.24$^{**}$ | 1.08 ± 0.03$^{ND}$ | 22.2 ± 1.75$^{ND}$ | 0.24 ± 0.012$^{ND}$ | 38.7 ± 1.80$^{*}$ | 1.65 ± 0.04$^{ND}$ | 7.3 ± 0.40$^{*}$ | 0.50 ± 0.011$^{ND}$ |
| ***Mock*** | *60.5 ± 2.29* | *1.13 ± 0.03* | *24.1 ± 1.77* | *0.23 ± 0.011* | *32.2 ± 1.81* | *1.66 ± 0.04* | *8.8 ± 0.59* | *0.51 ± 0.012* |
| **10 min** | 69.9 ± 2.18$^{ND}$ | 1.11 ± 0.04$^{ND}$ | 19.8 ± 1.63$^{**}$ | 0.26 ± 0.012$^{*}$ | 49.5 ± 1.85$^{**}$ | 1.45 ± 0.04$^{***}$ | 8.6 ± 0.55$^{ND}$ | 0.44 ± 0.010$^{*}$ |
| ***Mock*** | *64.7 ± 2.60* | *1.20 ± 0.05* | *28.4 ± 2.24* | *0.21 ± 0.013* | *42.1 ± 1.96* | *1.64 ± 0.04* | *9.8 ± 0.79* | *0.47 ± 0.012* |
| **20 min** | 72.1 ± 2.00$^{***}$ | 1.04 ± 0.04$^{ND}$ | 20.2 ± 1.55$^{***}$ | 0.23 ± 0.010$^{***}$ | 43.0 ± 1.41$^{*}$ | 1.56 ± 0.03$^{ND}$ | 6.2 ± 0.25$^{***}$ | 0.52 ± 0.007$^{***}$ |
| ***Mock*** | *60.4 ± 2.47* | *1.13 ± 0.04* | *34.6 ± 2.06* | *0.14 ± 0.007* | *37.9 ± 1.67* | *1.66 ± 0.04* | *13.4 ± 0.81* | *0.37 ± 0.009* |
| **30 min** | 78.9 ± 1.93$^{***}$ | 0.98 ± 0.03$^{**}$ | 17.1 ± 1.53$^{***}$ | 0.22 ± 0.010$^{ND}$ | 55.8 ± 1.76$^{***}$ | 1.50 ± 0.04$^{***}$ | 5.4 ± 0.41$^{***}$ | 0.59 ± 0.009$^{***}$ |
| ***Mock*** | *66.1 ± 2.24* | *1.13 ± 0.04* | *32.9 ± 2.35* | *0.20 ± 0.011* | *36.1 ± 1.58* | *1.72 ± 0.04* | *12.2 ± 0.74* | *0.44 ± 0.010* |
| **60 min** | 79.6 ± 1.99$^{ND}$ | 0.99 ± 0.04$^{*}$ | 21.6 ± 1.93$^{**}$ | 0.27 ± 0.014$^{***}$ | 56.0 ± 1.96$^{***}$ | 1.38 ± 0.04$^{***}$ | 6.2 ± 0.33$^{***}$ | 0.53 ± 0.010$^{**}$ |
| ***Mock*** | *76.4 ± 2.12* | *1.13 ± 0.04* | *31.3 ± 2.38* | *0.18 ± 0.008* | *45.5 ± 1.77* | *1.63 ± 0.04* | *10.1 ± 0.62* | *0.49 ± 0.010* |
| **Dose IAA** | **Density (%)** | **Skewness** | **Angle (°)** | **Parallelness** | **Density (%)** | **Skewness** | **Angle (°)** | **Parallelness** |
| ***Mock***[2] | *55.2 ± 1.67* | *1.28 ± 0.03* | *33.0 ± 1.35* | *0.22 ± 0.007* | *33.0 ± 1.22* | *1.81 ± 0.03* | *13.6 ± 0.54* | *0.31 ± 0.006* |
| **1 nM** | 70.9 ± 1.45$^{***}$ | 1.13 ± 0.03$^{***}$ | 25.5 ± 1.28$^{***}$ | 0.27 ± 0.010$^{***}$ | 44.7 ± 1.14$^{***}$ | 1.67 ± 0.03$^{***}$ | 10.1 ± 0.37$^{***}$ | 0.44 ± 0.007$^{***}$ |
| **10 nM** | 68.0 ± 1.42$^{***}$ | 1.19 ± 0.03$^{*}$ | 20.6 ± 1.08$^{***}$ | 0.24 ± 0.007$^{ND}$ | 43.4 ± 1.08$^{***}$ | 1.71 ± 0.02$^{*}$ | 8.6 ± 0.31$^{***}$ | 0.53 ± 0.006$^{***}$ |
| **100 nM** | 68.2 ± 1.47$^{***}$ | 1.19 ± 0.03$^{*}$ | 18.2 ± 0.95$^{***}$ | 0.30 ± 0.008$^{***}$ | 45.6 ± 1.03$^{***}$ | 1.71 ± 0.02$^{*}$ | 6.7 ± 0.19$^{***}$ | 0.58 ± 0.005$^{***}$ |
| **1 µM** | 62.9 ± 1.53$^{***}$ | 1.27 ± 0.03$^{ND}$ | 20.1 ± 1.01$^{***}$ | 0.29 ± 0.008$^{***}$ | 43.6 ± 1.16$^{***}$ | 1.66 ± 0.02$^{***}$ | 7.8 ± 0.29$^{***}$ | 0.51 ± 0.005$^{***}$ |
| **10 µM** | 62.9 ± 1.52$^{***}$ | 1.31 ± 0.03$^{ND}$ | 20.6 ± 1.00$^{***}$ | 0.31 ± 0.008$^{***}$ | 40.7 ± 1.10$^{***}$ | 1.80 ± 0.03$^{ND}$ | 8.9 ± 0.28$^{***}$ | 0.52 ± 0.006$^{***}$ |

[1]10 nM IAA treatments at various times. Density threshold 55.

[2]Dose series imaged at 20–30 min of treatment. Density threshold 45.

Values are ± standard error.

Time series, N = Approximately ≥ 10 cells (Region 2) and approx. 4–9 cells (Region 3) per root from 7 roots per treatment per timepoint.

Dose series, N = Approximately 8–12 cells per region per root; at least 10 roots per treatment..

*, $p \leq 0.05$; **, $p \leq 0.01$; ***, $p \leq 0.001$; $^{ND}$ = no difference; vs. mock, Student's t-test.

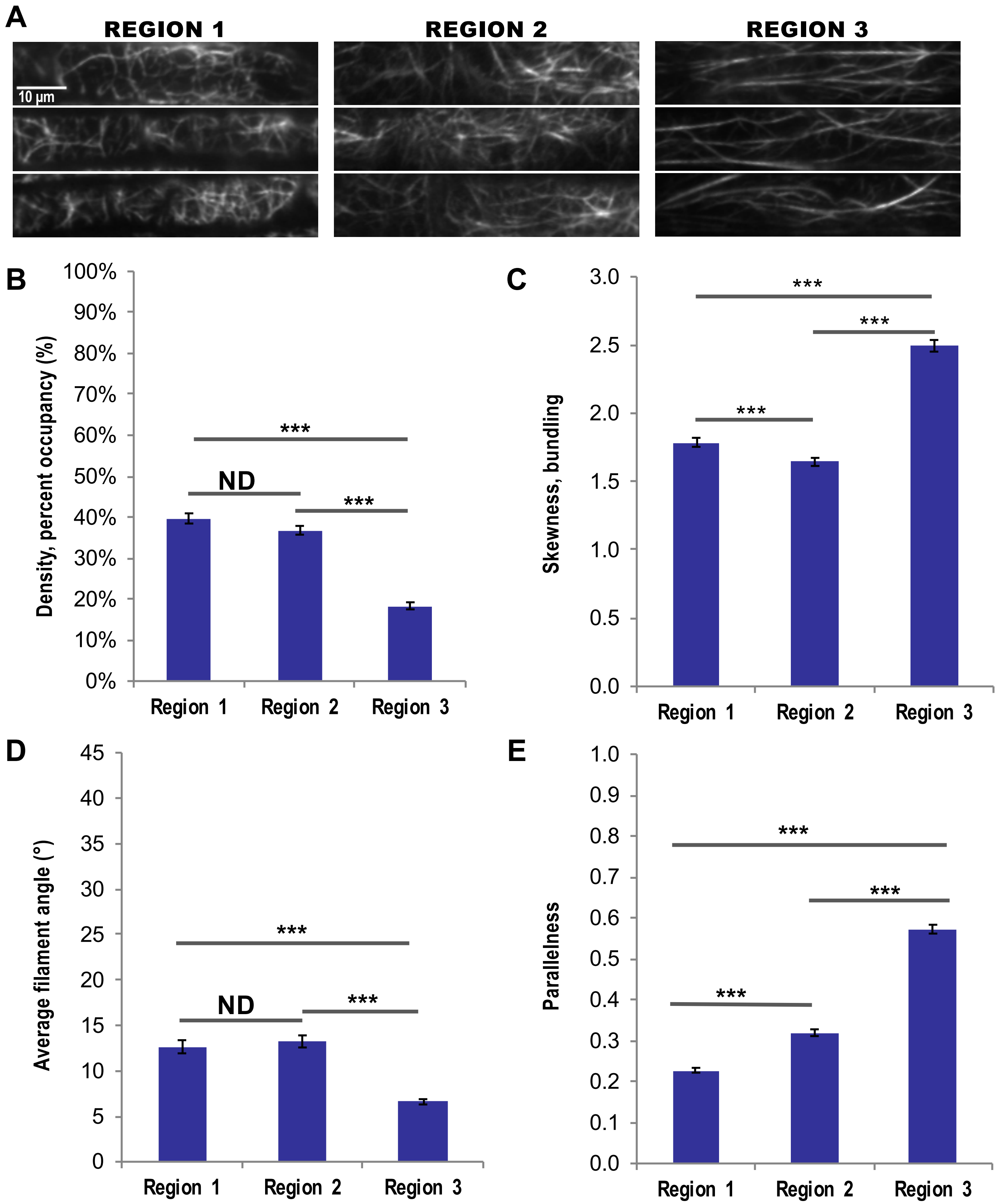

A
REGION 1
REGION 2
REGION 3
10 µm
B
Density, percent occupancy (%)
100%
90%
80%
70%
60%
50%
40%
30%
20%
10%
0%
***
ND
***
Region 1
Region 2
Region 3
C
Skewness, bundling
3.0
2.5
2.0
1.5
1.0
0.5
0.0
***
***
***
Region 1
Region 2
Region 3
D
Average filament angle (°)
45
40
35
30
25
20
15
10
5
0
***
ND
***
Region 1
Region 2
Region 3
E
Parallelness
1.0
0.9
0.8
0.7
0.6
0.5
0.4
0.3
0.2
0.1
0.0
***
***
***
Region 1
Region 2
Region 3

**Supplemental Figure 1.** Epidermal Cells in Different Root Regions Exhibit Distinct Actin Filament Arrays.

**(A)** Representative VAEM images of GFP-fABD2–labeled actin in epidermal cells from subjective root regions. Scale bar, 10 µm.

**(B)** to **(E)** Quantification of individual actin architecture or orientation metrics in three root regions. Results for density **(B)**, skewness **(C)**, angle **(D)**, and parallelness **(E)** showed significantly different actin arrays for most parameters, most notably between Regions 2 and 3. Actin filaments in Region 1 were dense and moderately bundled, with high average filament angle and low parallelness. Region 2 was characterized by a dense filament array, lower bundling, high average filament angle and filaments that were moderately parallel to each other. Region 3 was half as dense as Region 1 or 2, and exhibited a high degree of parallel, longitudinal bundles, with approximately 50% decrease in average filament angle and 40% increase in filament parallelness compared with Region 2.

N = 8–12 cells per region per root for 20 roots. ***$p \leq 0.001$; N.D., no differences; Student's t-test.

A

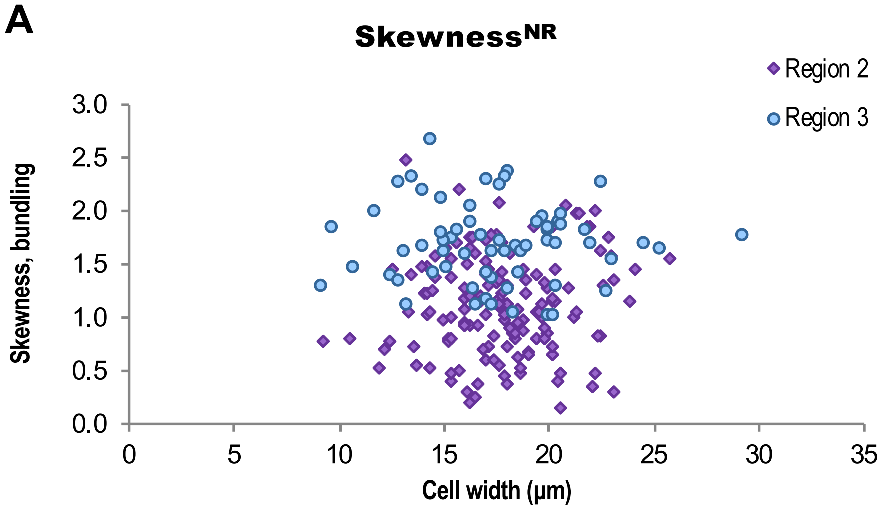

Skewness^NR
Region 2
Region 3
Skewness, bundling
3.0
2.5
2.0
1.5
1.0
0.5
0.0
0
5
10
15
20
25
30
35
Cell width (µm)


B

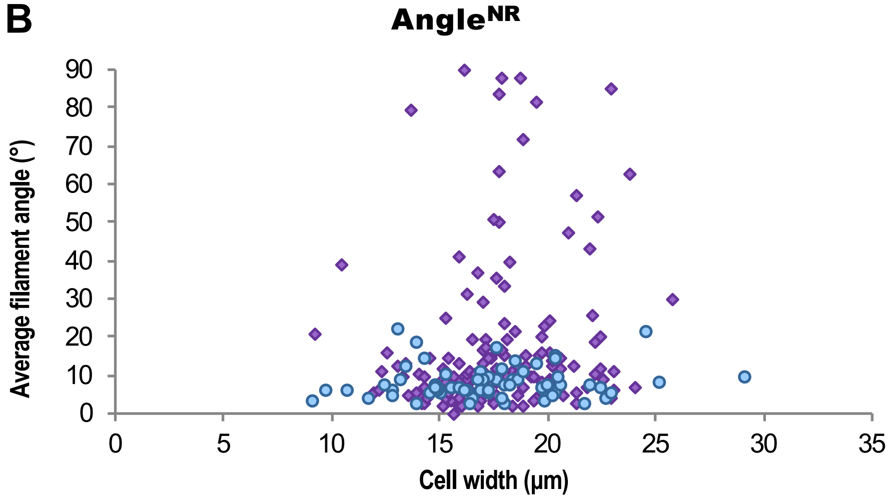

Angle^NR
Average filament angle (°)
90
80
70
60
50
40
30
20
10
0
0
5
10
15
20
25
30
35
Cell width (µm)


C

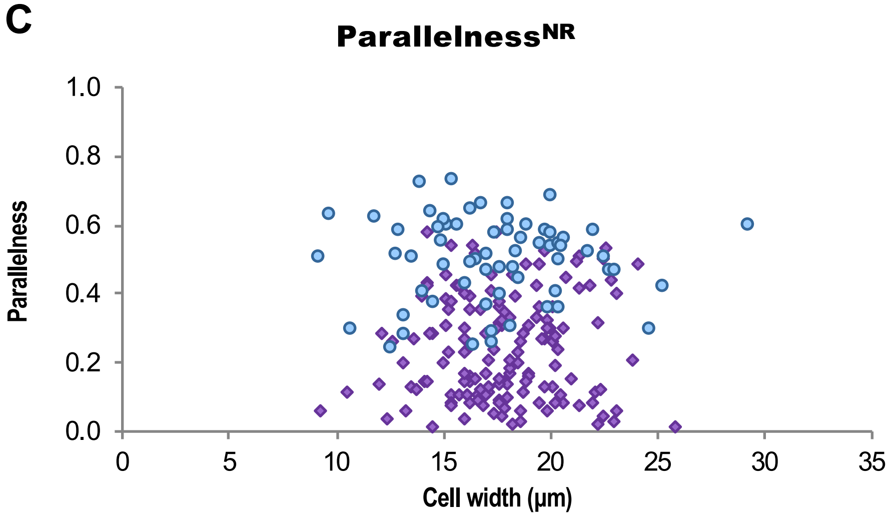

Parallelness^NR
Parallelness
1.0
0.8
0.6
0.4
0.2
0.0
0
5
10
15
20
25
30
35
Cell width (µm)

**Supplemental Figure 2.** Actin Filament Arrays Are Not Predictive of Cell Width.

**(A)** to **(C)** Quantification of individual actin architecture or orientation metrics plotted with respect to corresponding cell width in two root regions. Despite the high predictability between actin architecture parameters and cell length (shown in **Figure 1**) Filament architecture and orientation were not predictable based on cell width. Results for density **(Figure 1)** skewness **(A)**, angle **(B)**, and parallelness **(C)** vs. cell width showed no predictive relationships. Mean cell length, Region 2 = 57 ± 28 μm. Mean cell length, Region 3 = 128 ± 34 μm. Region 2 measurements are shown in purple diamonds; Region 3 in blue circles.

*Not shown*: mean actin filament density: Region 2 = 52.3 ± 0.02%; Region 3 = 15.4 ± 0.01%. Mean actin filament bundling/skewness: Region 2 = 1.12 ± 0.03; Region 3 = 1.71 ± 0.04. Mean filament angle: Region 2 = 17.5 ± 1.6°; Region 3 = 8.1 ± 0.5°. Mean filament parallelness: Region 2 = 0.24 ± 0.01; Region 3 = 0.50 ± 0.02.

N = 60–120 cells from 20 roots. NR, no predictive relationship, Bivariate fit/ANOVA. Results are from one experiment.

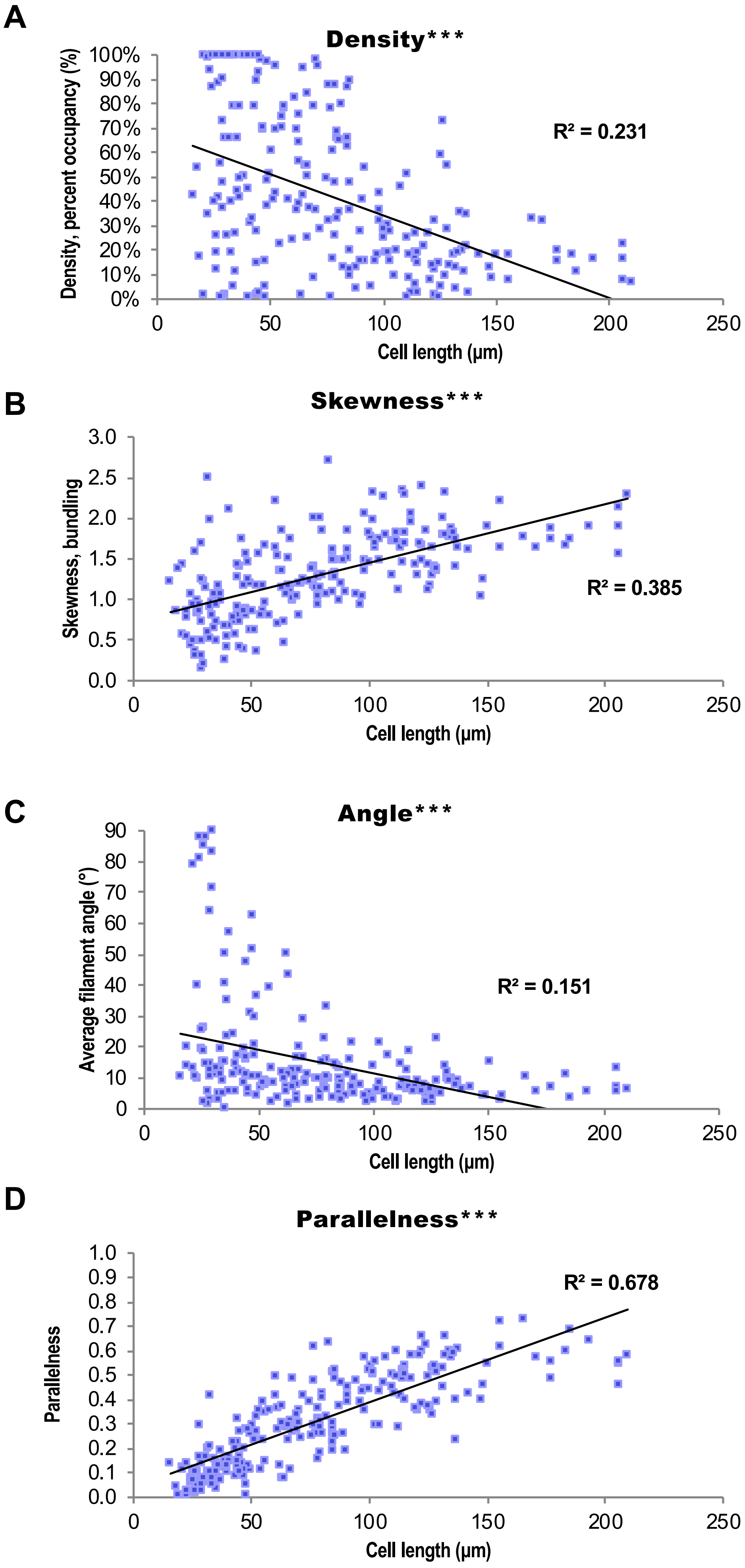
A
Density***
Density, percent occupancy (%)
100%
90%
80%
70%
60%
50%
40%
30%
20%
10%
0%
R² = 0.231
0
50
100
150
200
250
Cell length (μm)
B
Skewness***
Skewness, bundling
3.0
2.5
2.0
1.5
1.0
0.5
0.0
R² = 0.385
0
50
100
150
200
250
Cell length (μm)
C
Angle***
Average filament angle (°)
90
80
70
60
50
40
30
20
10
0
R² = 0.151
0
50
100
150
200
250
Cell length (μm)
D
Parallelness***
Parallelness
1.0
0.9
0.8
0.7
0.6
0.5
0.4
0.3
0.2
0.1
0.0
R² = 0.678
0
50
100
150
200
250
Cell length (μm)

**Supplemental Figure 3.** Actin Filament Arrays Are Predictive of Cell Length.

**(A)** to **(D)** Quantification of individual actin architecture or orientation metrics plotted with respect to corresponding cell length in two root regions, shown with $R^2$ values (linear fit determined in Excel) for reference only. These graphs are the same as **Figure 1** but are not shaded by region. Since it is not possible to know whether cell length or the aspects of actin organization is the independent variable, linear regression analysis is not an appropriate statistical metric on which to base strong conclusions.

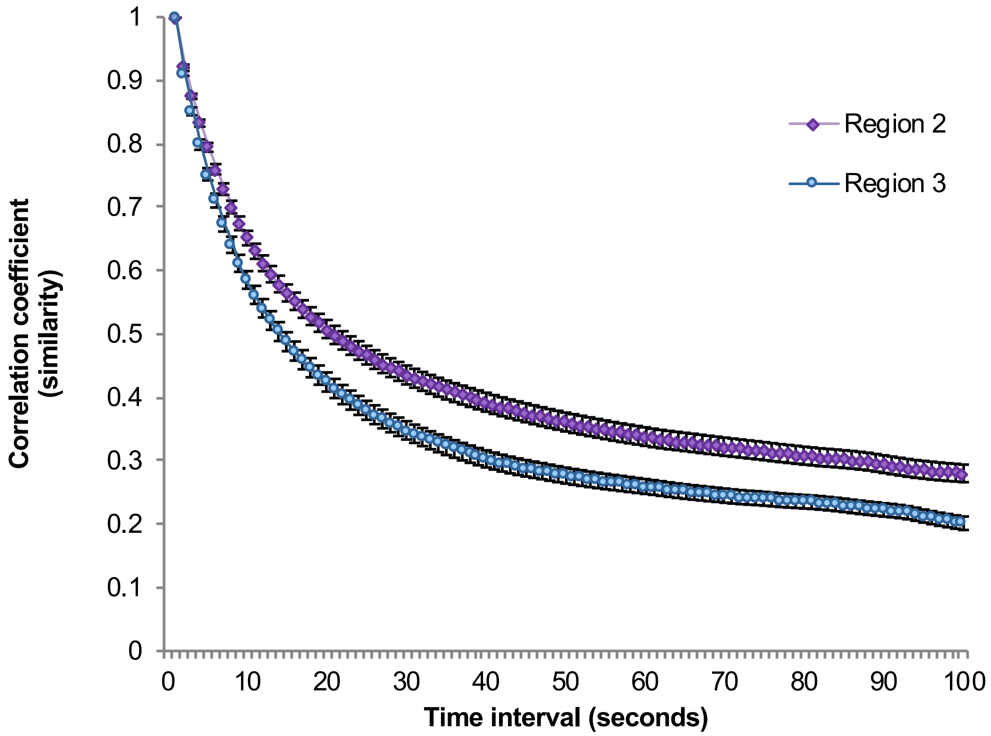


**Supplemental Figure 4.** Actin Arrays in Region 3 Are More Dynamic than in Region 2.

Filament arrays in short cells exhibited less overall dynamicity (i.e., short cells showed a slower decrease in similarity between pixel intensities and location between time intervals) compared with filament arrays of long cells.

100-s timelapse movies were collected from short and long cells in the same 30 roots. N = 80 (Region 3) to 148 (Region 2) cells from the same 30 plants as **Figure 2** and **Table 1**.  ***, F ≤ 0.001, one way ANOVA.

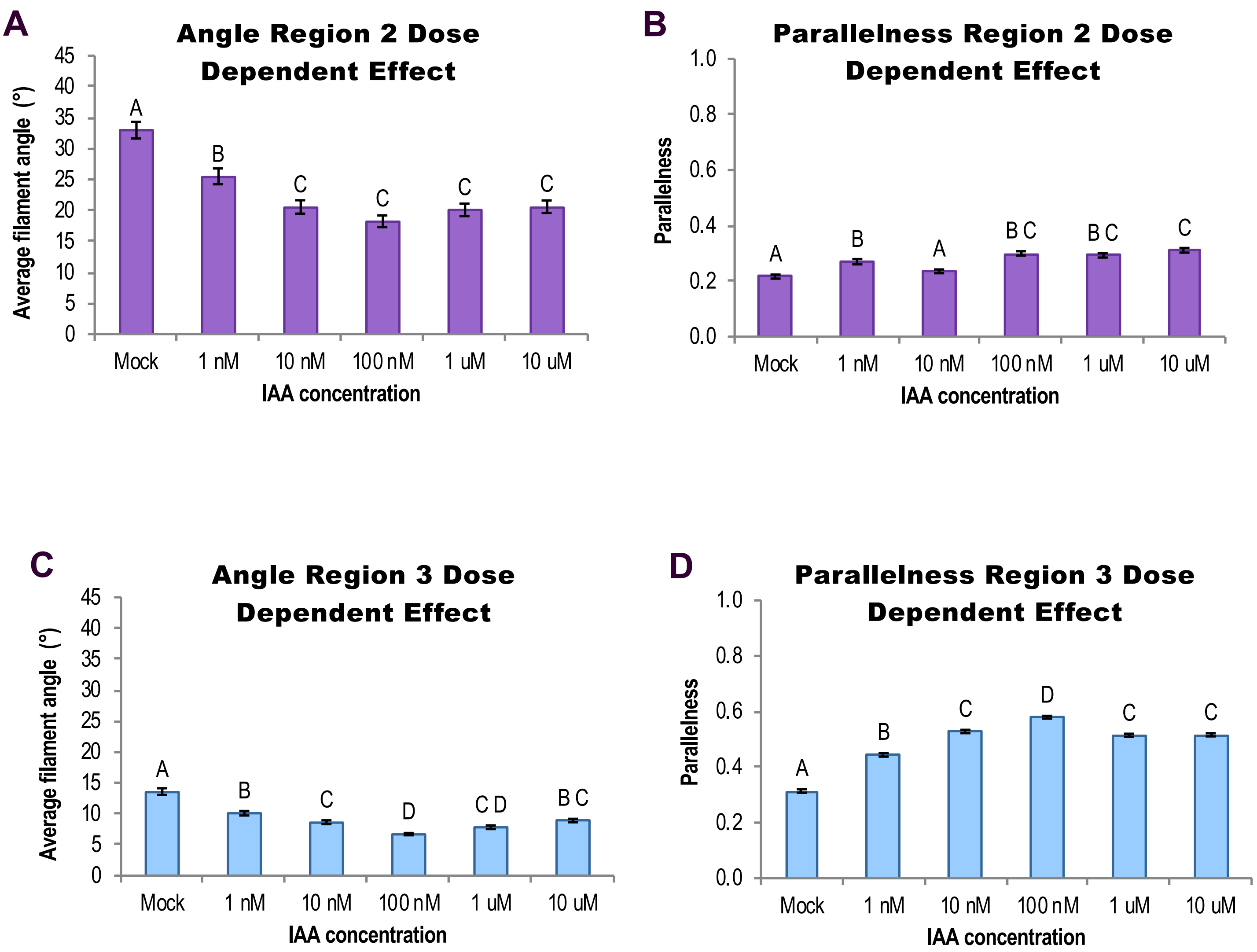


**Supplemental Figure 5.** Short-Term IAA Treatments Induce Dose-Dependent Changes in Actin Filament Organization.

**(A)** to **(D)** Quantification of actin orientation in root epidermal cells: IAA triggered a dose-dependent decrease in average filament angle **(A)** and **(C)** and increase in parallelness **(B)** and **(D)**. Region 2 measurements are shown in **(A)** and **(B)**; Region 3 in **(D)** and **(E)**. The dose-dependency was more pronounced in Region 3.

Cells whose lengths fell between 85 and 94 µm were counted in both regions. N = 8–12 cells per region per root from at least 10 roots per treatment. Different letters indicate statistically significant differences, oneway ANOVA, compared with Tukey-Kramer HSD in JMP (see Methods for more information). All IAA experiments were performed and analyzed double blind.

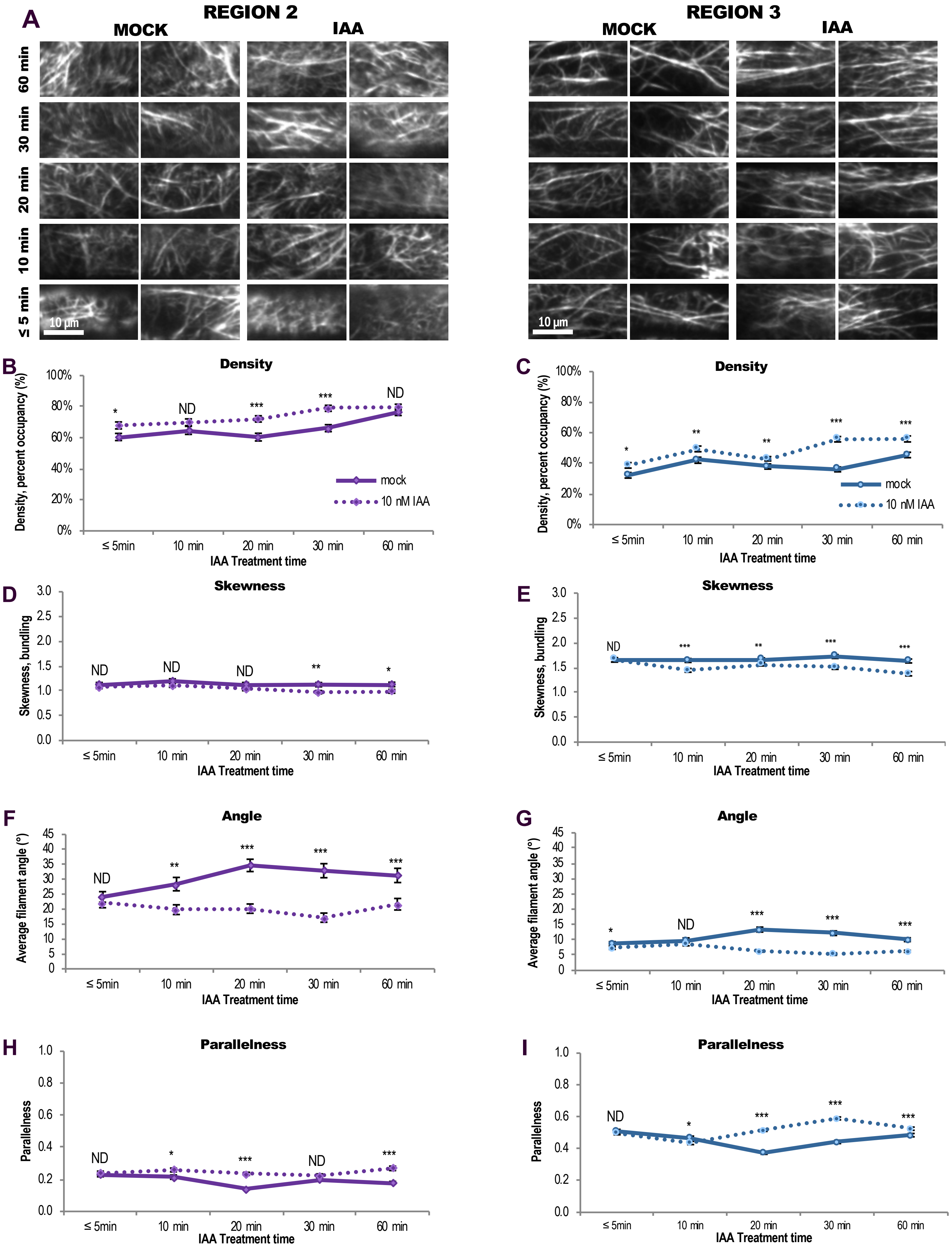
A
REGION 2
MOCK
IAA
REGION 3
MOCK
IAA
60 min
30 min
20 min
10 min
≤ 5 min
10 µm
10 µm
B
Density
Density, percent occupancy (%)
IAA Treatment time
mock
10 nM IAA
C
Density
Density, percent occupancy (%)
IAA Treatment time
mock
10 nM IAA
D
Skewness
Skewness, bundling
IAA Treatment time
E
Skewness
Skewness, bundling
IAA Treatment time
F
Angle
Average filament angle (°)
IAA Treatment time
G
Angle
Average filament angle (°)
IAA Treatment time
H
Parallelness
Parallelness
IAA Treatment time
I
Parallelness
Parallelness
IAA Treatment time
≤ 5min
10 min
20 min
30 min
60 min
ND

**Supplemental Figure 6.** Short-Term IAA Treatments Induce a Time-Dependent Increase in Actin Filament Density and Longitudinal Orientation.

**(A)** Representative VAEM images of GFP-fABD2–labeled actin in epidermal cells from Region 2 (left) and Region 3 (right). Scale bar, 10 μm.

**(B)** to **(E)** Quantification of actin architecture in Regions 2 and 3: 10 nM IAA (dotted lines) triggered a time-dependent increase in actin filament density **(B)** and **(C)** and decrease in skewness **(D)** and **(E)** compared to mock (unbroken lines). Region 2 measurements **(B)**, **(D)**, **(F)**, and **(H)** are shown in purple; Region 3 **(C)**, **(E)**, **(G)**, and **(I)** in blue.

**(F)** to **(I)** Quantification of actin orientation in Regions 2 and 3: after 10 nM IAA treatments, actin in both regions appeared more “organized”, with lower average filament angle **(F)** and **(G)** relative to the longitudinal axis of the cell and filaments generally more parallel to each other **(H)** and **(I)**. These responses were roughly time-dependent, with a peak in parallel longitudinality occurring after 20–30 min of treatment.

N = 7 roots per treatment per timepoint (≥ 10 cells from Region 2 and approx. 4–9 cells from Region 3 from each root). ND, no statistical differences; *, $p \leq 0.05$; **, $p \leq 0.01$; ***, $p \leq 0.001$, Student’s t-test of IAA (dotted line) vs. mock (unbroken line) on that region at that timepoint. Results are from one representative experiment of 2 similar experiments with similar results. All IAA experiments were performed and analyzed double blind.

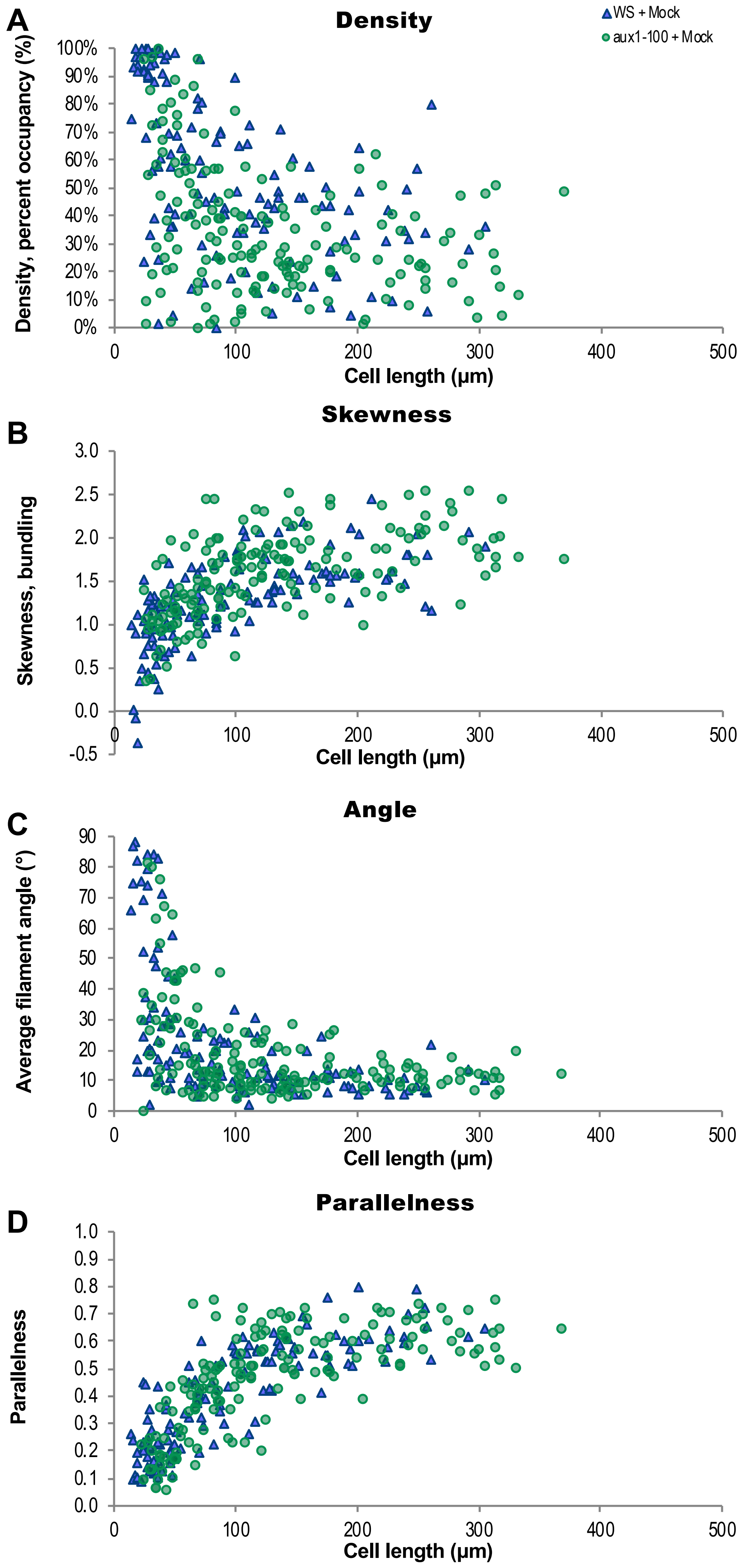
A
Density
WS + Mock
aux1-100 + Mock
Density, percent occupancy (%)
100%
90%
80%
70%
60%
50%
40%
30%
20%
10%
0%
0
100
200
300
400
500
Cell length (μm)
B
Skewness
Skewness, bundling
3.0
2.5
2.0
1.5
1.0
0.5
0.0
-0.5
Cell length (μm)
C
Angle
Average filament angle (°)
90
80
70
60
50
40
30
20
10
0
Cell length (μm)
D
Parallelness
Parallelness
1.0
0.9
0.8
0.7
0.6
0.5
0.4
0.3
0.2
0.1
0.0
Cell length (μm)

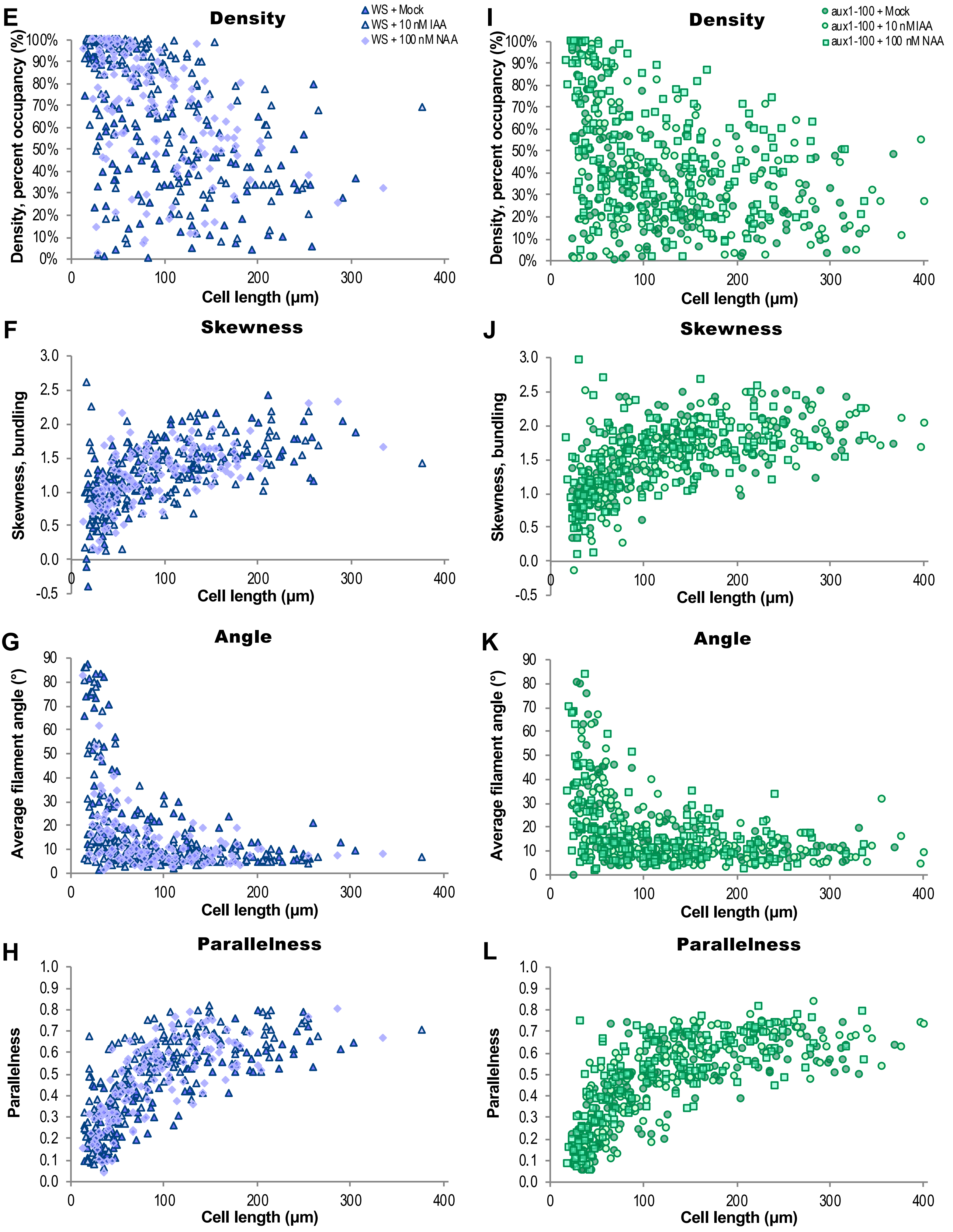
E
Density
WS + Mock
WS + 10 nM IAA
WS + 100 nM NAA
Density, percent occupancy (%)
Cell length (µm)
I
Density
aux1-100 + Mock
aux1-100 + 10 nM IAA
aux1-100 + 100 nM NAA
Density, percent occupancy (%)
Cell length (µm)
F
Skewness
Skewness, bundling
Cell length (µm)
J
Skewness
Skewness, bundling
Cell length (µm)
G
Angle
Average filament angle (°)
Cell length (µm)
K
Angle
Average filament angle (°)
Cell length (µm)
H
Parallelness
Parallelness
Cell length (µm)
L
Parallelness
Parallelness
Cell length (µm)

**Supplemental Figure 7.** Actin Filament Organization Plotted with Respect to Corresponding Cell Length in WS and *aux1-100*.

**(A)** to **(D)** Quantification of individual actin architecture or orientation metrics plotted with respect to corresponding cell length in WS and *aux1-100.*

**(E)** to **(H)** Quantification of WS individual actin architecture or orientation metrics after treatment with mock, IAA, and NAA, plotted with respect to corresponding cell length.

**(I)** to **(L)** Quantification of *aux1-100* individual actin architecture or orientation metrics after treatment with mock, IAA, and NAA, plotted with respect to corresponding cell length.

These graphs are scatterplots representing the same dataset as **Figure 5**; actin measurements were quantified on a per-cell basis and each data point represents a single cell's actin array plotted against its length.

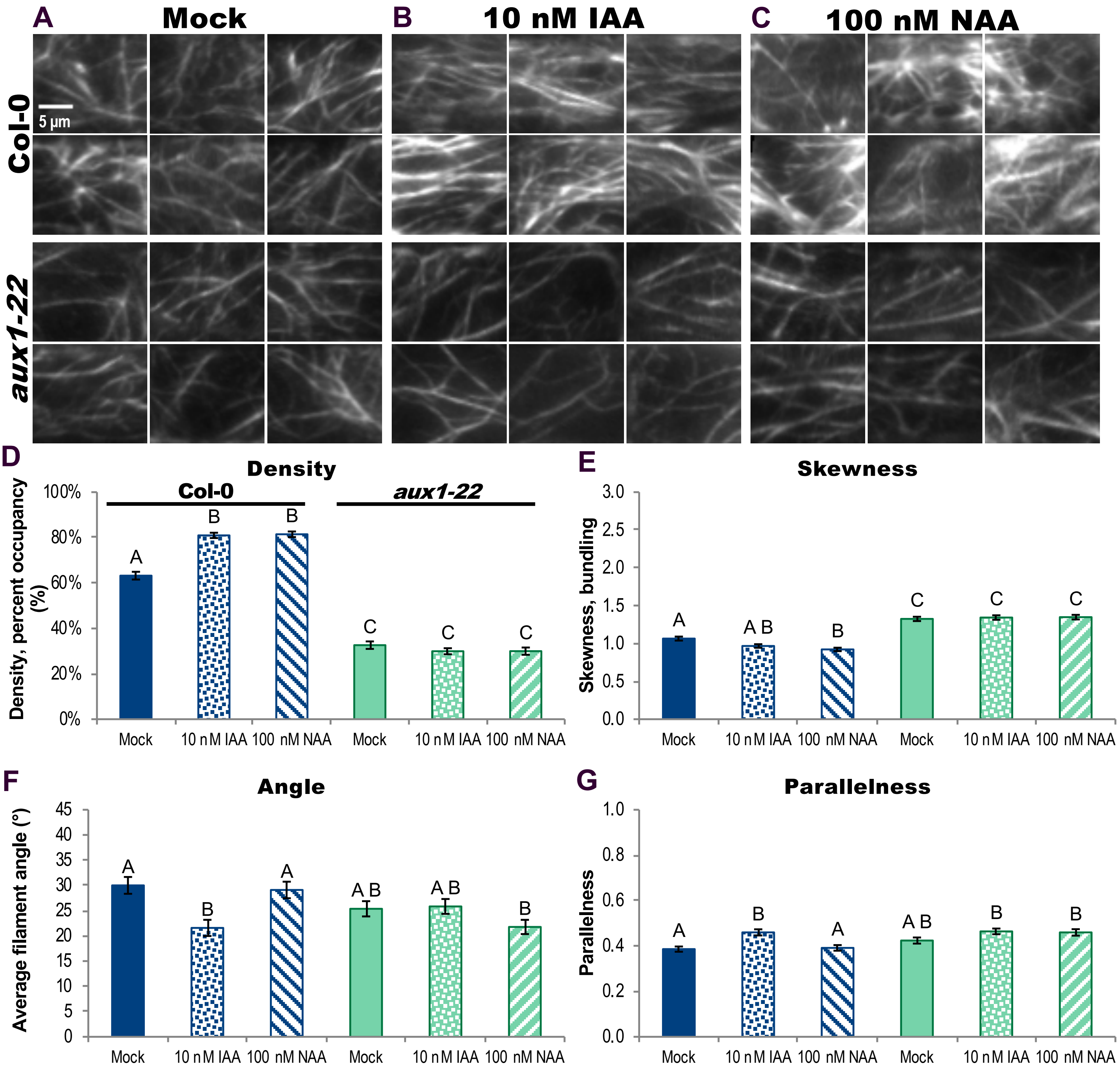

A
Mock
B
10 nM IAA
C
100 nM NAA
Col-0
5 µm
aux1-22
D
Density
Col-0
aux1-22
Density, percent occupancy (%)
100%
80%
60%
40%
20%
0%
A
B
B
C
C
C
Mock
10 nM IAA
100 nM NAA
Mock
10 nM IAA
100 nM NAA
E
Skewness
Skewness, bundling
3.0
2.5
2.0
1.5
1.0
0.5
0.0
A
A B
B
C
C
C
Mock
10 nM IAA
100 nM NAA
Mock
10 nM IAA
100 nM NAA
F
Angle
Average filament angle (°)
45
40
35
30
25
20
15
10
5
0
A
B
A
A B
A B
B
Mock
10 nM IAA
100 nM NAA
Mock
10 nM IAA
100 nM NAA
G
Parallelness
Parallelness
1.0
0.8
0.6
0.4
0.2
0.0
A
B
A
A B
B
B
Mock
10 nM IAA
100 nM NAA
Mock
10 nM IAA
100 nM NAA

**Supplemental Figure 8.** Actin Organization in *aux1-22* Fails to Respond to Short-Term IAA Treatments but Partially Responds to the Membrane-Permeable Auxin NAA.

**(A)** to **(C)** Representative VAEM images of GFP-fABD2–labeled actin in epidermal cells from wildtype (Col-0) and *aux1-22*, treated for 20–30 min with mock **(A)**, 10 nM IAA **(B)**, or 100 nM NAA **(C)**. Scale bar, 5 µm.

**(D)** to **(G)** Quantification of actin organization in root epidermal cells. Both IAA and NAA failed to trigger an increase in actin filament density **(D)** and decrease in skewness **(E)** in *aux1-22* but actin density in wildtype cells increased in response to both IAA and NAA and skewness decreased with both auxin treatments. Wildtype response is shown in blue and *aux1-22* in green; mock, solid; 10 nM IAA, dots; 100 nM NAA, stripes. After IAA treatment, actin arrays in wildtype plants were more "organized," with lower average filament angle **(F)** relative to the longitudinal axis of the cell and filaments generally more parallel to each other **(G)**. NAA triggered the increase in actin density in wildtype plants, but had no effect on angle or parallelness when measured on a per-cell basis. Average actin filament angle and parallelness in *aux1-22* failed to reorganize in response to IAA and only average filament angle decreased (and no increase in parallelness) with the membrane-permeable auxin NAA.

N = 5–38 cells per root; 10 roots per genotype per treatment. Different letters indicate statistically significant differences, oneway ANOVA, compared with Tukey-Kramer HSD in JMP (see Methods for more information). Actin measurements were quantified on a per-cell basis; see Methods for description and Supplemental for scatter plots. Results are from one experiment. All auxin experiments were performed and analyzed double blind.

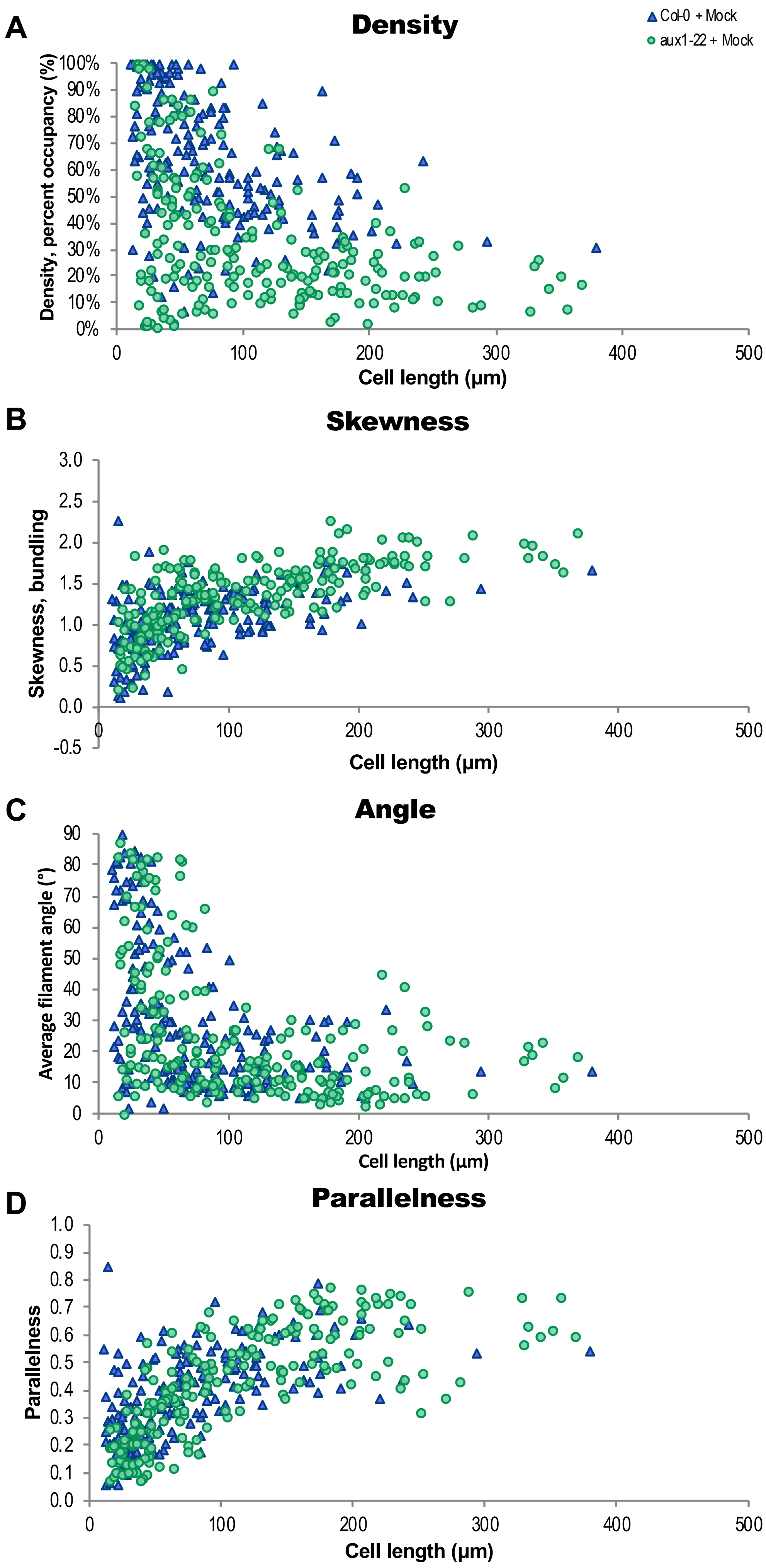
A
Density
Col-0 + Mock
aux1-22 + Mock
Density, percent occupancy (%)
100%
90%
80%
70%
60%
50%
40%
30%
20%
10%
0%
0
100
200
300
400
500
Cell length (μm)
B
Skewness
Skewness, bundling
3.0
2.5
2.0
1.5
1.0
0.5
0.0
-0.5
0
100
200
300
400
500
Cell length (μm)
C
Angle
Average filament angle (°)
90
80
70
60
50
40
30
20
10
0
0
100
200
300
400
500
Cell length (μm)
D
Parallelness
Parallelness
1.0
0.9
0.8
0.7
0.6
0.5
0.4
0.3
0.2
0.1
0.0
0
100
200
300
400
500
Cell length (μm)

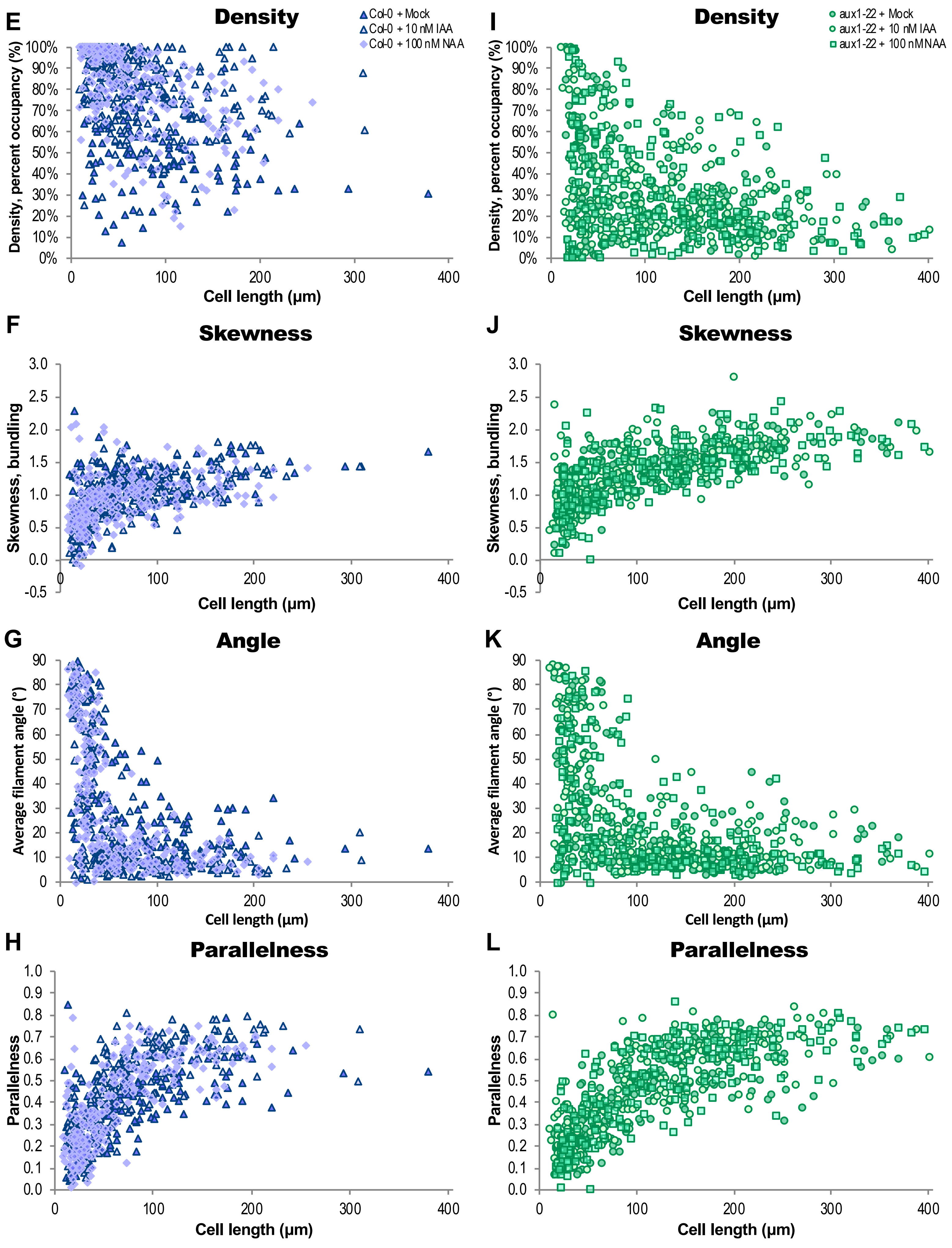

E
Density
Col-0 + Mock
Col-0 + 10 nM IAA
Col-0 + 100 nM NAA
Density, percent occupancy (%)
Cell length (µm)
I
Density
aux1-22 + Mock
aux1-22 + 10 nM IAA
aux1-22 + 100 nM NAA
Density, percent occupancy (%)
Cell length (µm)
F
Skewness
Skewness, bundling
Cell length (µm)
J
Skewness
Skewness, bundling
Cell length (µm)
G
Angle
Average filament angle (°)
Cell length (µm)
K
Angle
Average filament angle (°)
Cell length (µm)
H
Parallelness
Parallelness
Cell length (µm)
L
Parallelness
Parallelness
Cell length (µm)

**Supplemental Figure 9.** Actin Filament Organization Plotted with Respect to Corresponding Cell Length in Col-0 and *aux1-22*.

**(A)** to **(D)** Quantification of individual actin architecture or orientation metrics plotted with respect to corresponding cell length in Col-0 and *aux1-22.*

**(E)** to **(H)** Quantification of Col-0 individual actin architecture or orientation metrics after treatment with mock, IAA, and NAA, plotted with respect to corresponding cell length*.*

**(I)** to **(L)** Quantification of *aux1-22* individual actin architecture or orientation metrics after treatment with mock, IAA, and NAA, plotted with respect to corresponding cell length*.*

These graphs are scatterplots representing the same dataset as Supplemental Figure 8; actin measurements were quantified on a per-cell basis and each data point represents a single cell's actin array plotted against its length.

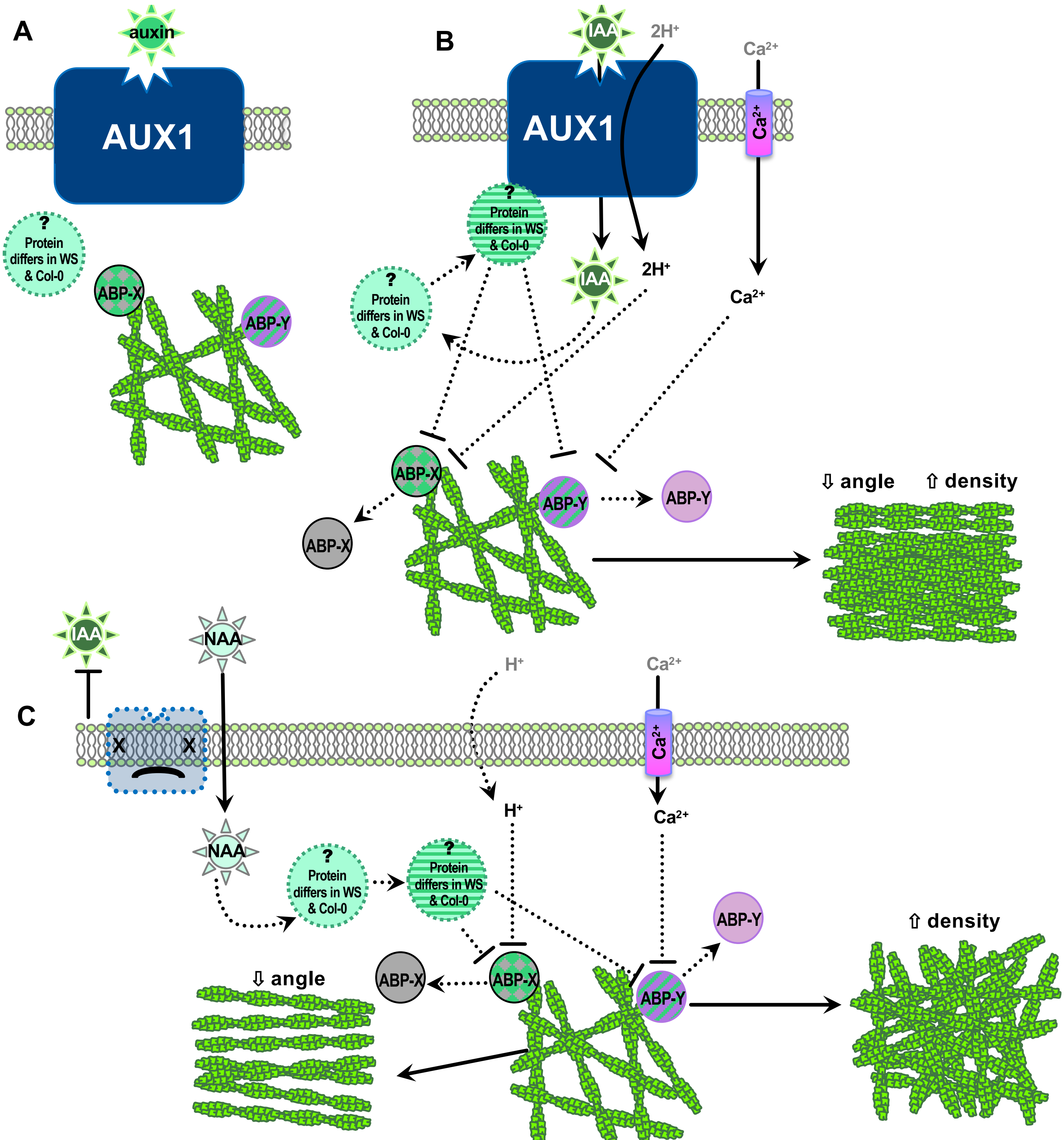


**Supplemental Figure 10.** Hypothetical model of auxin perception by AUX1 upstream of actin cytoskeleton reorganization.

**(A)** Control conditions: unidentified actin binding proteins (ABP) -X and -Y are active and maintain actin array; an unidentified intermediary that differs between Col-0 and WS is inactive.

**(B)** Auxin is transported into a cell by AUX1, activating the unknown intermediary, inactivating both actin binding proteins, and inducing increased actin filament abundance, decreased filament angle, and increased parallelness.

**(C)** In the absence of AUX1, the membrane permeable auxin NAA enters a cell, possibly activating the unknown intermediary; ABP-X and ABP-Y are differentially regulated and only either an increase in actin abundance or decrease in filament angle occurs.
